# Elovl1 inhibition reduced very long chain fatty acids in a mouse model of adrenoleukodystrophy

**DOI:** 10.1101/2024.11.11.622837

**Authors:** Jeremy Y. Huang, Brian Freed, Martin Hanus, Kelly Keefe, Ming Sum Ruby Chiang, Alexander Brezzani, Yongyi Luo, Yihang Li, Becky Lam, Stephanie Holley, Joseph Gans, Zuzana Dostalova, Buyun Tang, Clifford Phaneuf, Erin Teeple, Laura Parisi, Lilu Guo, Zhonglin Zhao, Sofia N. Kinton, Jacquelyn Dwyer, Sandrine Teixeira, Hong Ma, Gary Asmussen, Rajashree McLaren, Donghui Wang, Ann Baker, Craig S. Siegel, David Fink, Kristen Randall, Alexei Belenky, Suchitra Venugopal, Giorgio Gaglia, Jennifer Johnson, Dinesh S. Bangari, S. Pablo Sardi, Bailin Zhang, Alexander Michel, Jonathan D. Proto, Alla Kloss, Tatiana Gladysheva, Can Kayatekin

**Affiliations:** Sanofi

## Abstract

Adrenoleukodystrophy (ALD) is a rare neurometabolic disease caused by mutations in the *ABCD1* gene, which encodes for the peroxisomal very long chain fatty acid (VLCFAs) transporter. It is a debilitating disorder, which has a spectrum of clinical presentations. The most severe form is a rapidly progressing demyelinating disease called cerebral ALD or CALD. Patients with cALD have a life expectancy of 2-4 years after onset and symptoms often manifest in childhood. The other forms are adrenomyeloneuropathy or AMN, which is a slower progressing degeneration of the spinal cord, and adrenal insufficiency (Addison disease). Since the accumulation of VLCFAs are a common factor in all ALD pathologies, we identified therapeutic approach that could correct this metabolic defect. We developed a substrate reduction therapy (SRT) for ALD in the form of an inhibitor of the lipid elongase principally responsible for the generation of VLCFAs, Elovl1. This small molecule was able to successfully reduce the accumulation of VLCFA in the brain and spinal cord of *ABCD1*^−/y^ mice. We used single nuclei RNA seq to identify the pathways altered in the *ABCD1*^−/y^ mouse and corrected with Elovl1 inhibition. Though many lipid metabolism genes and pathways were indeed corrected, treatment with the Elovl1 inhibitor unexpectedly led to profound transcriptional changes beyond correction of pathways altered by loss of *ABCD1*. These data suggest that Elovl1 inhibition may have broader consequences in *ABCD1*^−/y^ mice than correction of lipid homeostasis.

## Introduction

Adrenoleukodystrophy, or ALD, is a rare neurometabolic disease that is caused by mutations in the ATP binding cassette D1 (*ABCD1)* gene.^1^ It has a spectrum of clinical presentations, which, in males, include adrenal insufficiency, a slowly progressing degeneration of the spinal cord (adrenomyeloneuropathy or AMN), or rapidly progressing demyelination of the brain (cerebral ALD or cALD) that can lead to death in 2-4 years after onset.^2,3^ Males can present with one or a combination of these three phenotypes. Women also suffer from ALD, despite carrying one functional copy of ABCD1. Due to the mosaic expression of either the dysfunctional or functional copy of ABCD1, females predominantly develop AMN; adrenal insufficiency and leukodystrophy are extremely rare in women.^4,5^ They also present with symptoms later in life than males and have slower disease progression. While there are no approved therapies for AMN, oral glucocorticoid treatment is used to treat adrenal insufficiency^6^ and hematopoietic stem cell transplants have demonstrated efficacy in halting the progression of cALD.^7–9^ More recently, ex-vivo lentiviral gene therapy has been approved for cALD, which may potentially allow patients who are ineligible for stem cell transplant or unable to find a donor to be treated.^10,11^ However, serious side effects of hematologic malignancies in a portion of the treated patients have been reported, resulting in FDA mandated black-box warnings^12^, which could prevent broad adoption and prophylactic treatment.

The *ABCD1* gene encodes for a peroxisomal membrane protein, called Abcd1 or ALDP (ALD-related protein)^13^ which plays a critical role in the degradation of very long chain fatty acids (VLCFAs). VLCFAs are imported into the peroxisome by Abcd1 as CoA-esters, where they are trimmed into long or short chain fatty acids.^14,15^ They are then transported to the mitochondria for further degradation. Hundreds of mutations in *ABCD1* have been discovered that impair the transport function of the protein, leading to accumulation of 24-carbon (C24) to 26-carbon (C26) VLCFAs in the plasma, brain, spinal cord, adrenal gland, and other tissues. It is this accumulated substrate, particularly the 26-carbon chain molecules, which are thought to impart a toxic effect that gives rise to the clinical manifestations of ALD. ^16,17^

In many monogenic rare diseases, substrate reduction therapy (SRT) has been used successfully in the clinic. For instance, Gaucher disease patients suffer from the loss of the enzyme glucocerebrosidase, which is responsible for the lysosomal degradation of glucosylceramide. Inhibiting the enzyme responsible for the synthesis of glucosylceramide, glucosylceramide synthase, is a highly effective approach that is FDA approved for the treatment of patients (Zavesca® (miglustat)^18^, Cerdelga® (eliglustat)).^19,20^ Similar approaches are currently in clinical trials for a variety of diseases, including Fabry disease (Venglustat ^21^), Niemann-Pick type-C disease (milglustat^22^) and Pompe disease (MZE001^23^). Beta-oxidation, *i.e.* the shortening of fatty acid chains, is opposed by fatty acid elongation, which is mediated by the Elovl family of enzymes. There are seven Elovl proteins in humans, each with specificity for certain carbon-chain lengths. The Elovl family member primarily responsible for the generation of fatty acids with carbon chains lengths of 22 to 26 is Elovl1.^24^ Thus, Elovl1 represents a promising SRT target for the treatment of ALD.

Here, we present a novel, orally bioavailable, brain penetrant Elovl1 inhibitor. This molecule was able to significantly reduce VLCFAs in the brain and spinal cord of mice deficient for Abcd1 function. To evaluate the benefit of this compound, we performed a biochemical and behavioral characterization of the *ABCD1^−/y^* mouse model. In the first 12 months of the life of this animal, there was little or no difference in rotarod performance out to 12 months of age. Moreover, we found a similar lack of difference in the open field test, which measures many motor and non-motor outcomes, and the Von-Frey test, which measures touch sensitivity of the footpads. These results indicated that the CNS and PNS were not significantly damaged by the presence of this mutation in the first year of the mouse’s life. Interestingly, our longitudinal biochemical analysis showed that the brain and more significantly the spinal cord progressively accumulated more and more C26:0 VLCFA throughout the lifespan of the animal. This motivated us to look more deeply into the transcriptional profiles of the cells within the CNS, to reveal more subtle signals for pathology. The results paint a mixed picture on the therapeutic potential of Elovl1 inhibition for the treatment of ALD.

## Results

### Elovl1 heterozygous deletion is well tolerated with significant changes of lipids in the CNS

Previous efforts to generate compounds targeting Elovl1 encountered dose limiting toxicities in several animal species, including rodents.^25,26^ Moreover, Elovl1 knockout mice display neonatal lethality due to water loss caused by impaired epidermal barrier formation.^27^ To credential the safety of the target, we generated heterozygous Elovl1 knockout animals. CRISPR-Cas9 was used to delete 1958 bp of the *ELOVL1* gene, removing exons 2-8. The effect of this deletion was confirmed by qPCR, which showed that the relative level of *ELOVL1* transcripts in the brains of *ELOVL1^−/+^* mice were 59% of the WT animals, in line with the expected 50% reduction (Figure 1A). The loss of one copy of *ELOVL1* affected the C26:0 levels of tissues in proportion to the amount of C26:0 that naturally accumulated in that tissue. In plasma and liver, where there is normally very little C26:0, *ELOVL1^+/-^* were statistically indistinguishable from their WT counterparts (Figure 1B). In the kidneys, which have higher levels of C26:0, heterozygous animals had roughly half the amount of C26:0 as WT animals. In brain and spinal cord, where C26:0 levels are the highest, the differences were the most significant. *ELOVL1^+/-^* animals lost nearly all the C26:0 that would normally be present in these tissues. We performed a histopathological analysis of a broad panel of tissues and found no abnormalities at 6 months of age, suggesting that even the large changes in lipids seen in the CNS were well tolerated (Figure 1C).

**Figure 1:**
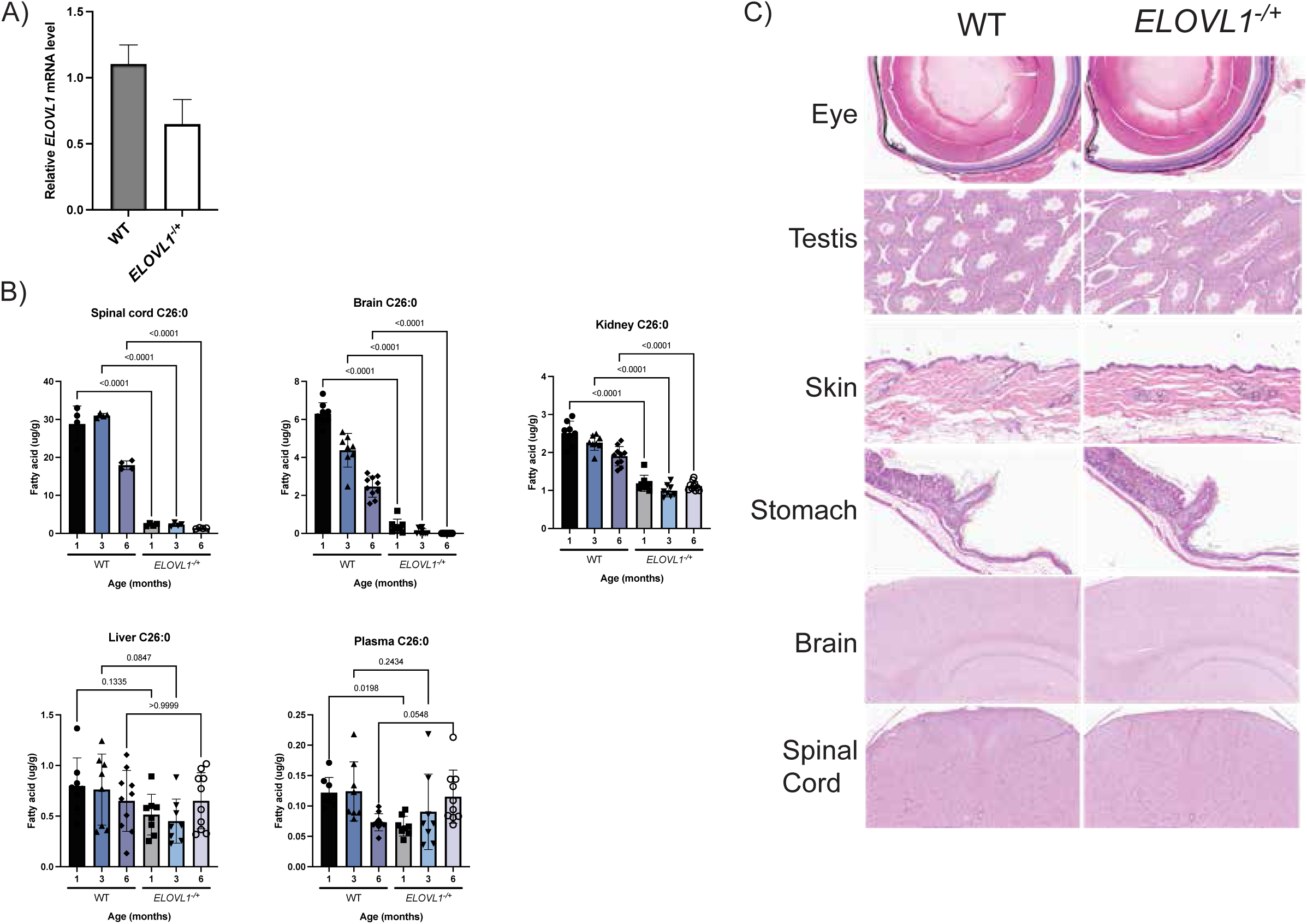
**A)** qPCR analysis of relative *ELOVL1* expression in brains of WT and heterozygous *ELOVL1* knockout mice (2 WT, 4 mutant). **B)** Total C26:0 fatty acid quantification as a function of mouse age across tissues (Spinal cord n = 4, Brain, Kidney, Liver n = 8-10, mixed sex). **C)** Representative images from H&E-stained tissue sections.

### CPD37 is a potent, selective inhibitor of Elovl1

To identify novel Elovl1 inhibitors, we screened ∼725,000 compounds using an *in vitro* microsomal assay for Elovl1 activity. Development of the initial hits from this screen led to the identification of CPD37 (Figure 2A). To confirm activity with intact cells, we measured the potency of CPD37 in reducing the amount of sphingomyelin, an abundant sphingolipid, bearing a C26:0 fatty acid chain. Fibroblasts isolated from childhood cerebral adrenoleukodystrophy (CCALD) patients were treated with compound for 72 hours at 37 °C and the ratio of SM C26:0 to C22:0 was measured by mass spectrometry. The molecule was highly effective, reducing the pool of lipids with an EC_50_ of 52 nM (Figure 2B). Treatment of patient fibroblasts with compound and ^13^C-C22 labeled substrate followed by GC-MS analysis of the intact cells confirmed that CPD37 also lowers total C26:0 fatty acid levels (Figure 2C).

**Figure 2:**
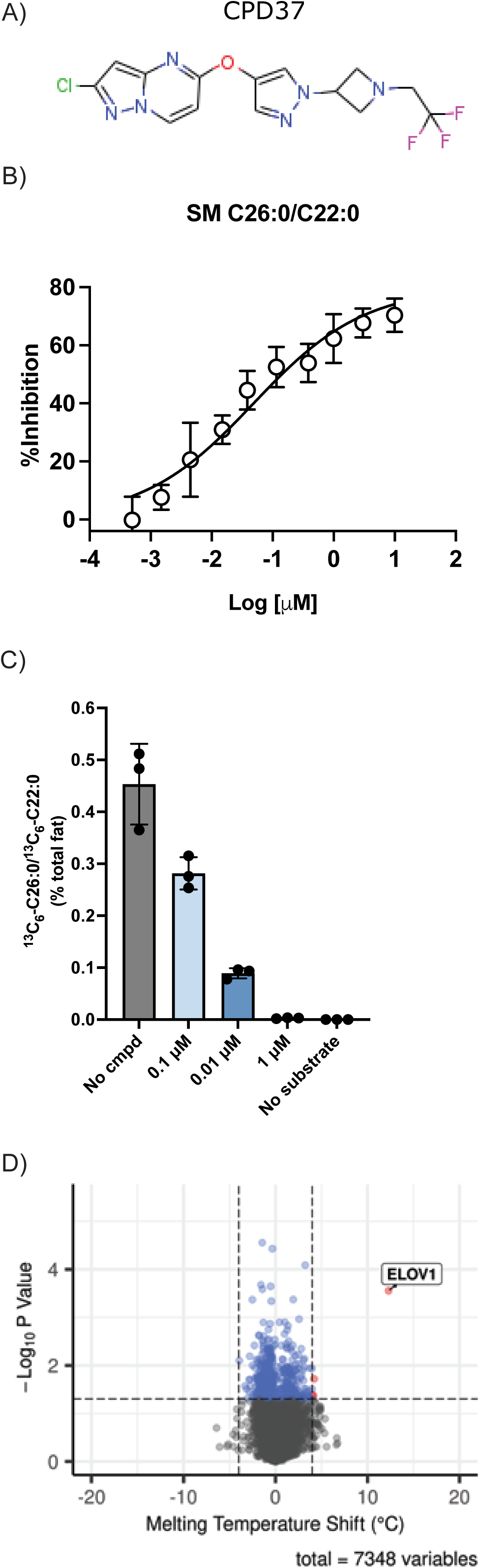
**A)** Structure of CPD37. **B)** Inhibition of sphingomyelin (SM) C26:0 upon treatment of ALD-patient fibroblasts with CPD37, normalized to SM C22:0. **C)** Inhibition of elongation of a stable-isotope labeled substrate with CPD37 treatment. **D)** Cellular thermal shift assay (CETSA) results from intact Jurkat cells treated with and without CPD37.

To examine the selectivity of CPD37, we performed a cellular thermal shift assay (CETSA). This method analyzes the melting point of proteins in the presence or absence of a compound to determine the targets of that compound. If a molecule binds a protein, the melting point of that protein is typically shifted to higher temperatures. By performing this assay in cell lysates, it is possible to determine the specificity of a compound by evaluating its impact on the melting point of all detectable proteins. Intact Jurkat cells were treated with compound for one hour and then lysates were analyzed at 8 temperatures by LC-MS. Most proteins detected were not shifted in their melting profiles, while Elovl1 was the most prominently shifted protein with a ΔT_m_ of 12.3 °C (Figure 2D). These results supported the hypothesis that CPD37 is quite specific for Elovl1. Further support for the selectivity of CPD37 was demonstrated by the absence of off target activity in selected profiling assays at 10 μM. In targeted enzymatic assays, CPD37 was also highly selective for Elovl1 over Elovl3 and Elovl7 (Table 1).

| physicochemical properties and CNS distribution |  | plasma protein binding<sup>d</sup> |  |
| --- | --- | --- | --- |
| Log D (pH 7.4) | 2.46 | human PPB (% bound) | 88.3 |
| Caco2 P<sub>Total</sub> A2B (10<sup>-7</sup> cm/s)<sup>a</sup> | 618 | mouse PPB (% bound) | 86.7 |
| efflux ratio<sup>b</sup> | 1.7 | rat PPB (% bound) | 88.8 |
| mouse brain/plasma K<sub>p</sub><sup>c</sup> | 1.10 |  |  |

| selectivity profiling |  |  |  |
| --- | --- | --- | --- |
| kinase selectivity (>50% inhibition at 10 μM) | 0/60 | hERG automated patch clamp IC<sub>50</sub> (μM) | >30 |
| off-target selectivity (>50% inhibition at 10 μM)<sup>e</sup> | 0/74 | hCaV1.2 IC<sub>50</sub> (μM) | >30 |
| HepG2 DILI IC<sub>50</sub> (μM) / Hep cytotoxicity IC<sub>50</sub> (μM) | >512 / >512 | ELOVL7 IC<sub>50</sub> (μM) | >99 |
|  |  | ELOVL3 IC<sub>50</sub> (μM) | >10 |

<sup>a</sup>LC/MS quantification of apical/basolateral transport at 20 μM in Caco2/TC7 cell line <sup>b</sup>Ratio of LC/MS quantification of B to A over A to B at 1 μM in Caco2/TC7 cell line <sup>c</sup>Brain/plasma partition coefficient determined from exposure in C57BL/6 mice at 2 hr after 3 mg/kg PO dosing <sup>d</sup>PPB measured at 5 μM <sup>e</sup>Binding, inhibition and uptake assays performed at Eurofins

### CPD37 is brain penetrant and reduced VLCFA in mice

Based on the promising *in vitro* activity and selectivity of CPD37, we performed PK/PD studies in mice to evaluate its properties *in vivo*. The pharmacokinetic parameters are summarized in table 1. Orally administered CPD37 had a bioavailability of 46%, reached a maximum blood concentration after 30 minutes, and was cleared with a half-life of 1.8 hours (Figure 3A). The exposure was proportional to the dose administered. To measure the blood-brain-barrier penetration of the compound, mice were intravenously infused with 3 mg/kg of CPD37 and exposure was measured at 4 time points in both the blood and the brain (Figure 3B). The exposure in both tissues paralleled each other and the exposure ratio between the brain and plasma was 1.1 to 1 (Table 1). WT mice were administered compound once per day (QD) by oral gavage for 12 days and the amount of C26:0 in the brain and spinal cord was measured to evaluate the efficacy of Elovl1 inhibition. Although the total C26:0 levels were numerically lower than vehicle treated animals at all doses, only the changes in the 30 mg/kg/day and 100 mg/kg/day treated animals reached statistical significance (Figure 3C). Since Elovl1 inhibition would be expected to block elongation beyond C20, we also examined the pools of C20:0 fatty acids. These FA should be elevated upon Elovl1 inhibition. This measurement was less sensitive to Elovl1 inhibition than C26:0 changes, as only the highest treatment dose showed a statistically significant increase in the spinal cord (Figure 3D).

**Figure 3:**
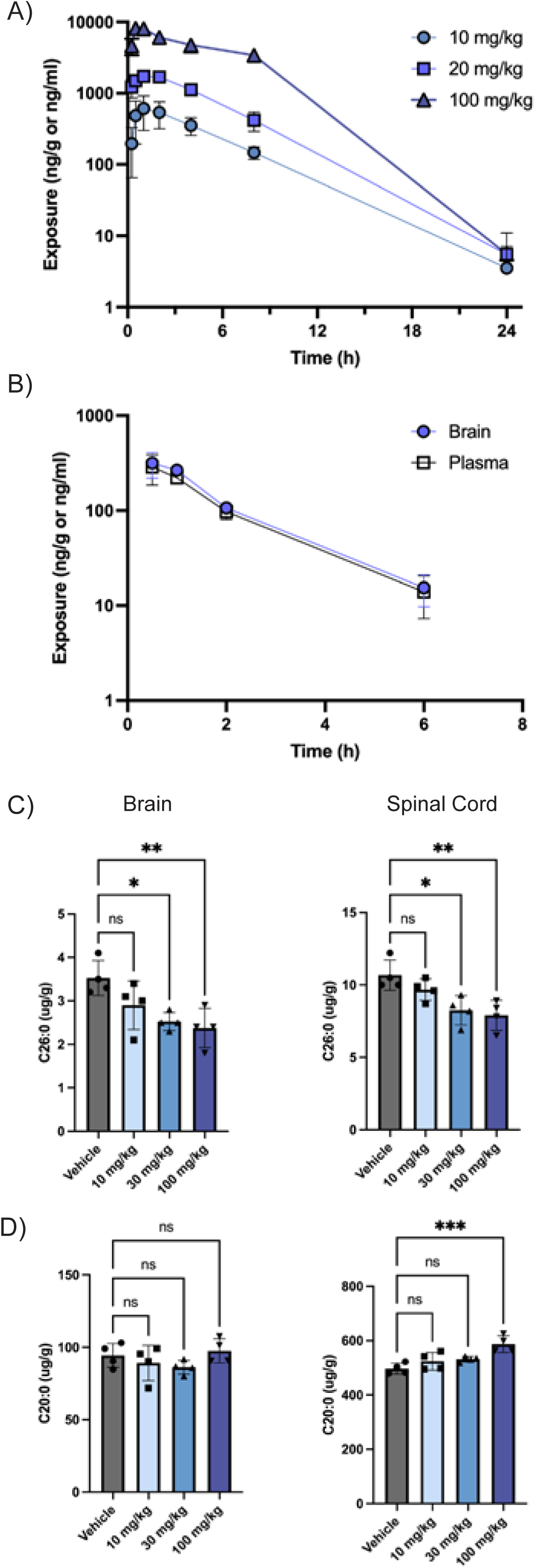
**A)** Plasma exposure of CPD37 following a single oral administration in mice (N=3, FVB, males). **B)** Plasma and brain exposure of CPD37 following a 3 mg/kg IV administration (N=3, FVB, males). **C)** Total C26:0 levels in the brain and spinal cord of WT mice treated with CPD37 for 12 days (N=4, FVB, males). **D)** Total C22:0 levels in the brain and spinal cord of WT mice treated with CPD37 for 12 days (N=4, FVB, males).

### Stable isotope labeled elongation assay in vivo

A major challenge in measuring a pharmacodynamic effect with CPD37 in mice was the slow turnover of VLCFA in the CNS. Even daily treatment with high doses of Elovl1 inhibitor for 12 days had modest effects on the total levels of C26:0 (Figure 3C), making it difficult to establish a dose response. We sought to overcome this challenge by directly measuring *de novo* elongation *in vivo* using stable isotope labeled metabolic tracing. For three days, P17-P20 pups were treated with ^13^C-labeled stearic acid (C18:0) followed by CPD37 1 hour later. This allowed for measuring the elongation of ^13^C-labeled FAs from C20:0 to C22:0 and beyond. In all cases, we attempted to measure elongation to C26:0 but sensitivity limitations often prevented detection beyond C22:0 or C24:0; C26:0 species were never detectable. Here, a dose-dependent change in the elongation was more apparent, not only in the increased accumulation of ^13^C-C20:0 (Figure 4A, left), but also in the decreased accumulation of ^13^C-C22:0 (Figure 4A, right). Notably, the changes in the labeled pools were statistically significant while the change in the total pool was not (Figure 4B), an indication of the increased sensitivity of this technique. All tissues evaluated (brain, spinal cord, liver) showed dose dependent changes in FA elongation (Figure 4). In tissues with higher amounts of lipid synthesis like the liver, elongation to ^13^C-C24:0 could also be detected (Figure 4D), which were even more impacted by Elovl1 inhibition.

**Figure 4:**
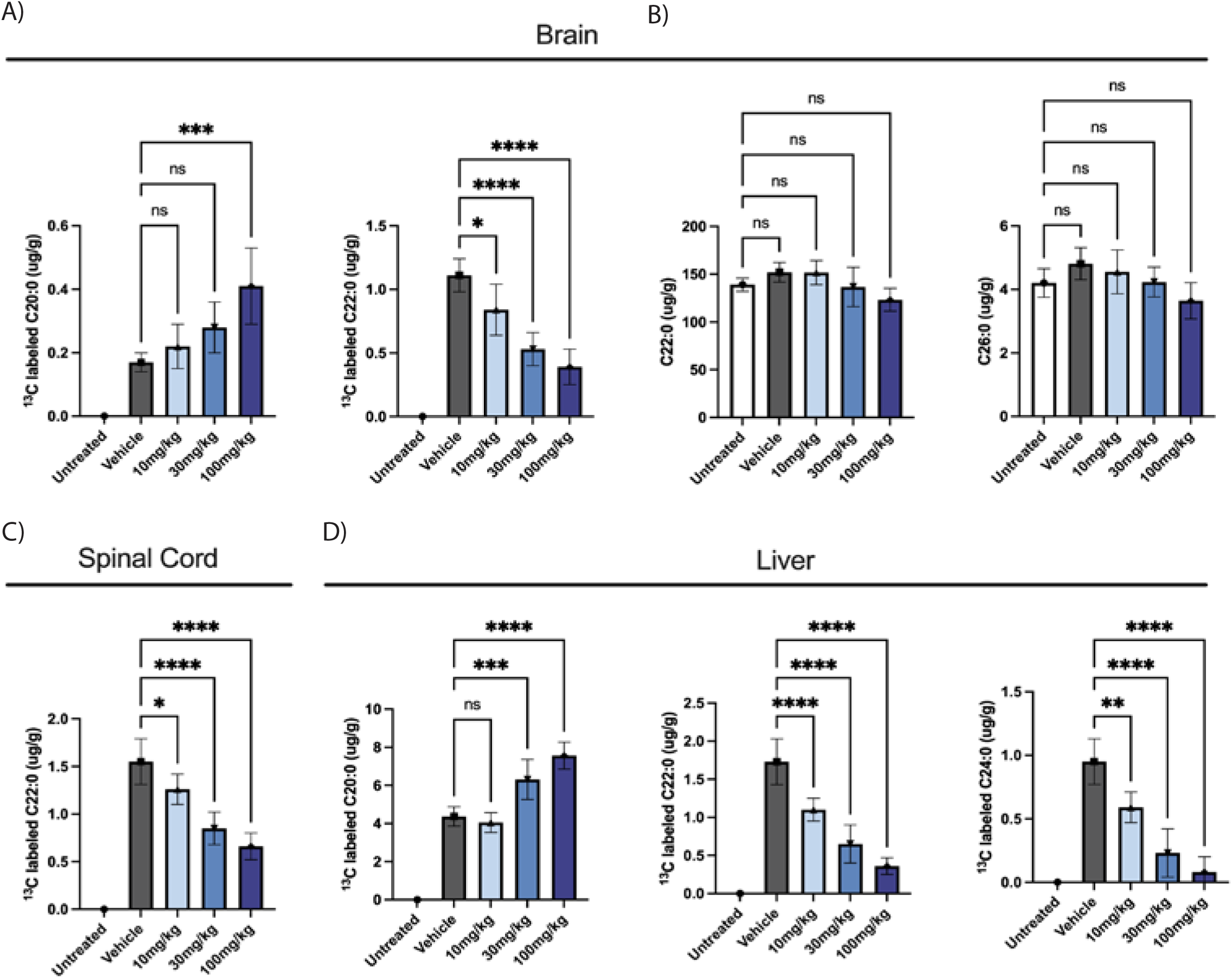
Stable isotope labeled *in vivo* elongation assay in WT mice. **A)** ^13^C-labeled C20:0 and C22:0 levels in the brain of mice treated with CPD37. **B)** Total C22:0 and C26:0 levels. **C)** Levels of ^13^C-labeled C22:0 in the spinal cord. **D)** ^13^C-labeled C20:0, C22:0, C24:0 levels in the liver. Vehicle (2%PVP k25, 0.1%SLS in water) n = 5, 10 mg/kg n = 6, 30 mg/kg n = 10, 100 mg/kg n = 9, mixed sex.

### ABCD1 model characterization

We performed a longitudinal analysis of total FA content in the brain, spinal cord, kidney, and liver of *ABCD1*^−/y^ and WT mice ^28–30^. The lipid levels at 1 week of age were taken as the baseline and log2-fold changes from the baseline are shown at 2, 4, 8, 16, 32, 52, and 78 weeks of age in mutant and WT mice (Figure 5). In the brain and spinal cord, patterns of accumulation were similar between *ABCD1^−/y^* and WT mice. Many of the FA with carbon chains shorter than 18 were either decreased or constant following birth, while many of the longer chain FAs steadily increased throughout the lifespan of the animal. In the brain, C22:0 and C24:0 VLCFAs showed different patterns of accumulation. In WT animals, there was a peak early in life, followed by reduction into adulthood, after which the levels remained constant. In *ABCD1^−/y^* animals, the FA levels rose into adulthood and, for some lipids, started to come down much later in life. In the brain and spinal cord, C26:0 levels showed the most dramatic difference between the two genotypes. Whereas the WT animals had a small increase into adulthood, followed by constant levels, *ABCD1^−/y^* animals continued to accumulate this FA throughout their life. This pattern of accumulation was especially prominent in the spinal cord, which also had the highest absolute levels of this FA.

**Figure 5:**
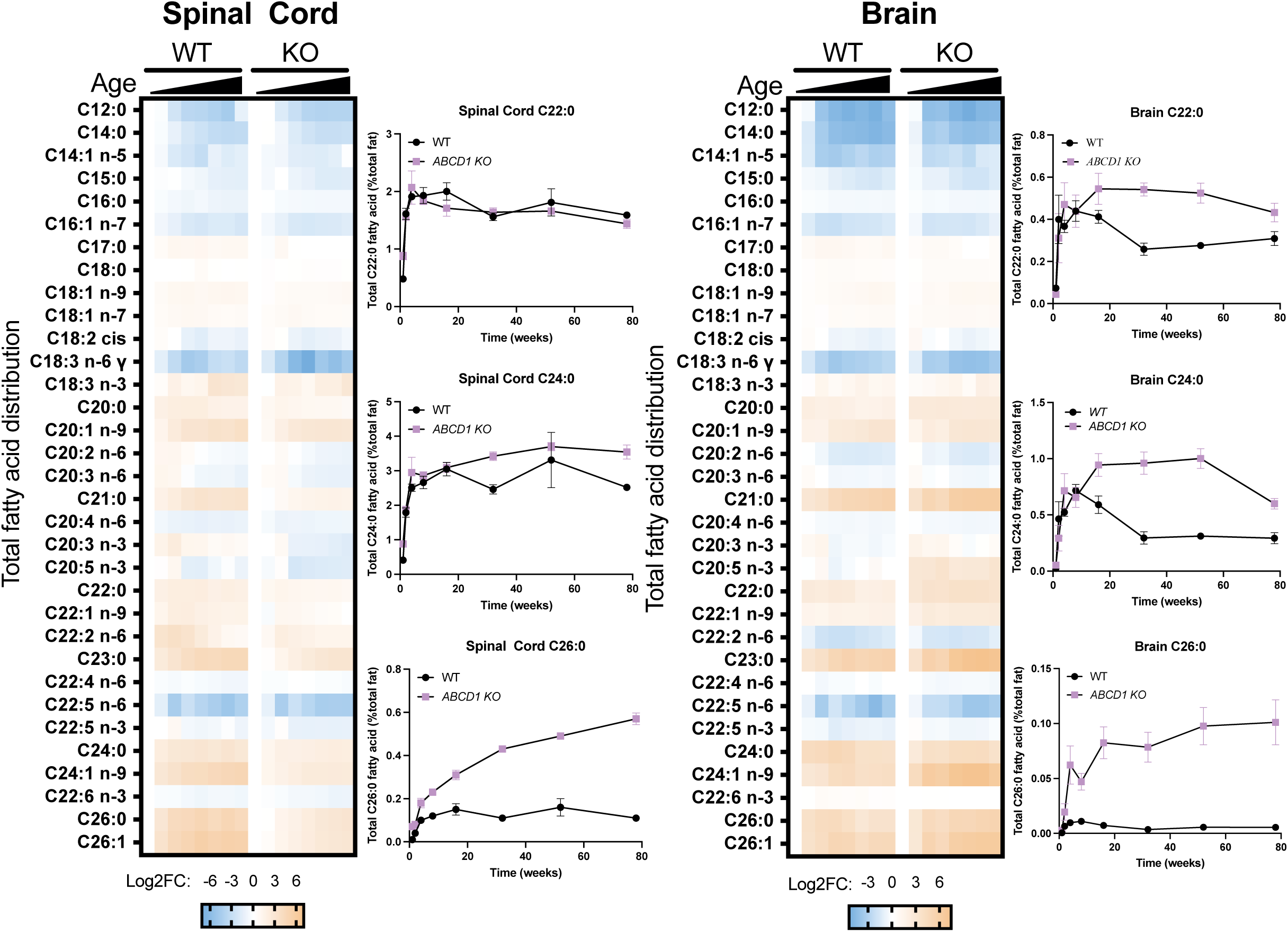
Changes in total fatty acids of various chain lengths and saturation states as a function of mouse age. The lipid levels at 1 week of age were taken as the baseline and relative changes are shown at 2, 4, 8, 16, 32, 52, and 78 weeks of age. N = 5 for all mutant ages, n = 3 for WT 1 week, n = 4 for WT 52 weeks, n = 5 for all other WT ages.

With the notable exception of C26:0 in the spinal cord, most lipids seemed to peak within the first 12 months of the animal’s life (Figure 5). Behavioral studies in these mice have previously only found differences very late in life.^31^ We sought to identify behavioral differences in the time span that could be used for an early efficacy determination. We performed a longitudinal behavioral analysis of male *ABCD1*^−/y^ mice, evaluating their performance on rotarod, Von Frey sensitivity, and the open field test. No differences were measurable in the in the open field test outcomes, including distance traveled and speed of movement (Figure S1A). In addition, there were no consistent, age-dependent differences in the foot pad sensitivity between WT and KO animals as measured by the Von Frey test (Figure S1B), or in rotarod performance (Figure S1C).

### Elovl1 inhibition reduced VLCFA levels in *ABCD1^−/y^* animals

Having biochemically and behaviorally characterized the *ABCD1*^−/y^ animals, we sought to reduce VLCFA levels *in vivo* using Elovl1 inhibition. 3-month-old mice were treated daily with orally administered CPD37 and VLCFA levels were measured in plasma, brain, and spinal cord following 7, 14, and 30 days of treatment. Plasma levels of C26:0 were reduced in a dose dependent manner, reaching WT-like levels at the highest dose of 100 mg/kg/day (Figure 6A, left). Plasma exposures were consistent with a linear dose dependence with no accumulation with repeated administration (Figure 6B). There was little time dependence to the reduction of VLCFA, with levels remaining roughly stable beyond 7 days of treatment. To our surprise, levels of C26:0 in the brain and spinal cord were little changed even after 30 days of treatment at the highest dose (Figure 6A, right). This is in part due to additional accumulation of VLCFA during the timeframe of the study. In the vehicle group, there was a time dependent increase in VLCFA in the brain. In the treatment groups, brain VLCFA remained constant, suggesting the beneficial effect at this young age may prevent further accumulation.

**Figure 6:**
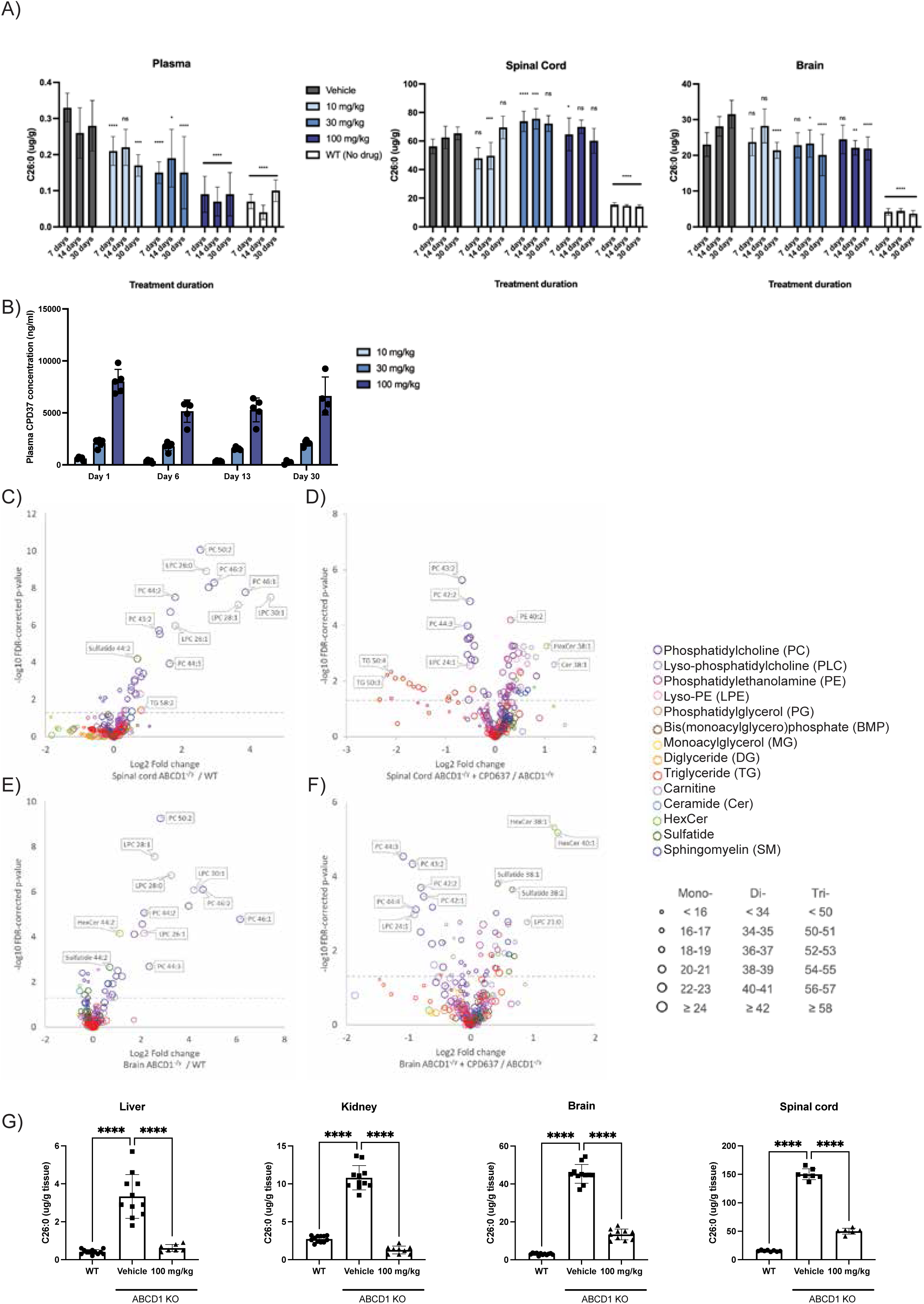
**A)** Changes in total C26:0 levels in *ABCD1^−/y^* mice following daily oral administration of CPD37 at three doses: 10 mg/kg; 30 mg/kg and 100 mg/kg for 7, 14, and 30 days (n = 7-8 male animals). Statistics are shown relative to the untreated age-matched ABCD1^−/y^ mice. Two-way ANOVA followed by Dunnett’s multiple comparisons test. **** indicates adjusted p-value <0.0001, *** indicates adjusted p-value <0.001, ** indicates adjusted p-value <0.01, * indicates adjusted p-value <0.05, ns = not significant. **B)** Plasma exposure of CPD37 measured 2 hours post dose (n = 5 animals) **C-F)** Untargeted lipidomics showing accumulation of PCs with ≥42 carbons were reduced with CPD37 treatment. Volcano plot for identified lipids, colored by lipid class and sized according to total number of carbons in monoacyl-, diacyl- and triacyl-chains. Dashed line indicates 0.05 FDR-corrected p-value. Full data are available Table S1. **C)** Spinal cord lipids in *ABCD1^−/y^* vs. WT mice. **D)** Spinal cord lipids in *ABCD1^−/y^* vs. CPD37-treated *ABCD1^−/y^* mice. **E)** Brain lipids in *ABCD1^−/y^* vs. WT mice. **F)** Brain lipids in *ABCD1^−/y^* vs. CPD37-treated *ABCD1^−/y^* mice. **G)** Changes in C26:0 levels after 7-month treatment of *ABCD1*^−/y^ mice with 100 mg/kg/day of CPD37 formulated in feed (n = 10-12 male animals).

The lack of more prominent changes after 30 days of treatment led us to explore the kinetics of lipid turnover following Elovl1 inhibition. Therefore, we performed untargeted lipidomic profiling to examine the effects of CPD37 on individual lipids in the brain and spinal cord (Figure 6C-F). We found that the accumulation of specific lipids in the *ABCD1^−/y^* mice, in particular PC lipids with ≥42 carbons, was more noticeably reduced by CPD37 (Figure 6C and E). Moreover, lowering was most significant for phosphatidylcholines, compared to other lipid classes such as sphingomyelins and ceramides (Figure 6D and F). There were notable reductions in some triglycerides in the spinal cord of treated animals, but these were restricted to a few lipids, rather than the entire class (Figure 6D).

To better evaluate the potential of our compound to reduce VLCFA in the CNS, we treated 6-month-old animals for 7 months with CPD37 formulated to deliver 100 mg/kg/day in feed. With these older animals, much of the accumulation of VLCFA has already happened. Combined with the longer treatment, we expected to see a more significant reduction of VLCFA in the brain and spinal cord. Indeed, this was the case. VLCFA levels were at WT levels in the kidney and below WT levels in the liver (Figure 6B, left). In the brain and spinal cord, VLCFA levels were lower by 70% and 66% respectively (Figure 6B, right).

### Single nuclei sequencing analysis of *ABCD1*^−/y^ mice and the effects of CPD37

To better characterize and understand long-term cell type dependent effects of *ABCD1^−/y^* and CPD37 administration, we performed single nuclei RNA sequencing on the three groups of animals from the long-term feed treatment study described above: the WT group, the vehicle treated *ABCD1^−/y^* group, and the *ABCD1^−/y^* + 100 mg/kg CPD37 compound. Brain and spinal cords from four mice per group were harvested following treatment, preprocessed (as described previously^32^), and sequenced (Figure 7A). After cell typing using the gene-set based algorithm SARGENT^33^ with cell type definitions and marker references^34^, we were able to robustly identify (Figures S2A and S2B) glial cells (Astrocytes, Astrocyte-Restricted Precursors, Microglia, OPCs, and Oligodendrocytes), supporting cells (Arachnoid barrier, Choroid plexus epithelium, and Vascular cells), and neural subtypes (Neural stem, GABA, Glutamatergic, Cholinergic, and Dopaminergic neurons) across the three treatment groups (Figures 7B and 7C).

**Figure 7:**
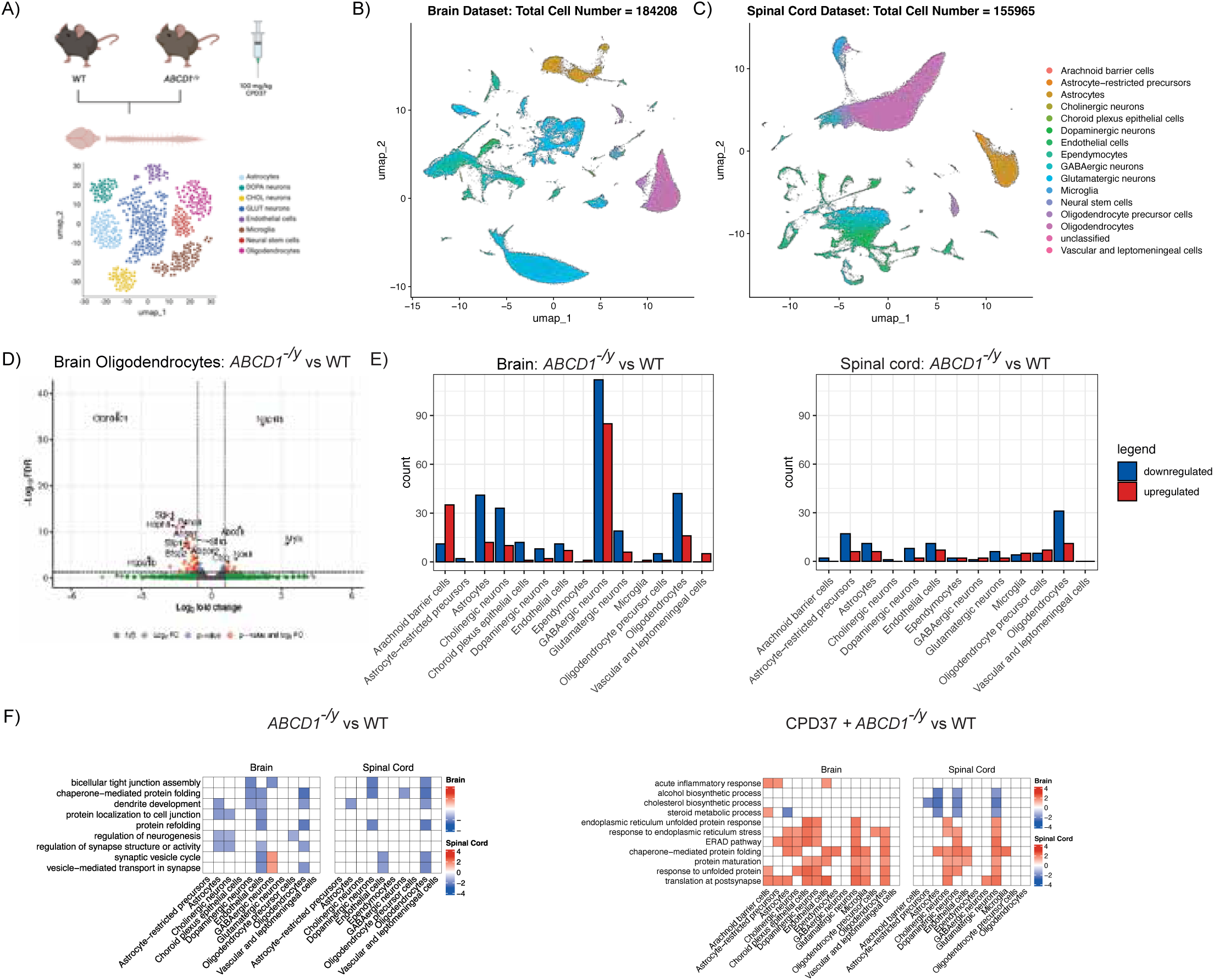
**A)** Single-Cell RNA-seq study design of effects of *ABCD1^−/y^* and effects of Elovl1 inhibitor CPD37 in mouse brain and spinal cord. Generated in Biorender.com **B)** UMAP visualization of all cells from twelve samples from the mouse brain cell-type annotated by SARGENT. **C)** UMAP visualization of all cells from twelve samples from the mouse spinal cord cell-type annotated by SARGENT **D)** (top) Volcano plot of differentially expressed genes by pseudobulk differential expression comparison in Mouse Oligodendrocytes in the Brain comparing *ABCD1^−/y^* model with Wild-Type mice. **E)** Grouped bar plot comparing number of genes which are up or downregulated in a DGE comparison between *ABCD1^−/y^* model and Wild-Type mice across cell types in (left) brain and (right) spinal cord. **F)** Heatmap of gene ontology pathway NES score from genes comparison by cell type, split by tissue in (left) *ABCD1^−/y^* model and Wild-Type mice and (right) CPD37 + *ABCD1^−/y^* model and *ABCD1^−/y^* model comparisons.

We observed that *ABCD1^−/y^* had limited effects on overall proportion of cell types except for nominal decreases in glutamatergic neurons (p-value = 0.012) in the spinal cord (Figures S3A and S3B). CPD37 treatment had similar limited effects. These observations were confirmed using a cell type-agnostic KNN-based method to assess differential abundance, Milo.^35^ No statistically significant cellular neighborhoods (transcriptionally similar groups of cells) were found to be differentially abundant in *ABCD1^−/y^* vs Wild-Type comparisons (Figure S4A). A limited group of cellular neighborhoods, retroactively associated with GABAergic neurons, were enriched in the CPD37 + *ABCD1^−/y^* treatment group (Figure S4B) in comparison to the *ABCD1^−/y^* + vehicle group (Figure S4C), yet the overall proportions of cellular neighborhoods were balanced across all cell types in both the brain and spinal cord.

Since effects of mutation status and treatment status were found to be cell type abundance agnostic, we performed pseudobulk differential gene expression analysis to ascertain potential pathway-level effects of *ABCD1^−/y^* and CPD37 treatment. In the *ABCD1^−/y^* mouse model, ABCD1 transcript elevation was observed across cell types, including oligodendrocytes (Figure 7D) as compared to WT controls, an expected compensatory mechanism for *ABCD1^−/y^*. CPD37 administration had a profound effect on *ABCD1^−/y^* mouse (Figures 7E and S5A). While many genes which were dysregulated by Abcd1 loss-of-function returned to Wild-Type expression levels (Figure S5B), including VLCFA-associated ^36^ protein folding pathway (Figures 7F, left and S5C) genes such as *AHSA1*, *CHORDC1*, *STIP1*, *FKPB4*, and many other genes and pathways were engaged post-treatment in comparison to untreated *ABCD1^−/y^* (Figure 7F, right) and WT (Figure S5C). Taken together, CPD37 administration normalized some *ABCD1^−/y^* instigated VLCFA-linked aberrant protein folding and transport functions in cell types such as oligodendrocytes and endothelial cells but resulted in increased stress responses in the brain (Figure S5D) and downregulated sterol processes in neurons and astrocytes in both the brain and the spinal cord.

## Discussion

In this study, we investigated the potential for Elovl1 inhibition to treat the underlying pathological feature of adrenoleukodystrophy (ALD), the accumulation of very long chain fatty acids (VLCFA). To better understand the safety liabilities of this target, we generated heterozygous *ELOVL1* knockout animals, which revealed tissue-specific reductions in C26:0 levels. These reductions were most prominent in the CNS and less prominent in the periphery. Histopathological assessments indicated no adverse effects at six months, suggesting that partial loss of Elovl1 function and the concomitant reduction of VLCFAs was well tolerated. Convinced of the apparent safety of reducing Elovl1 function, we discovered a new, oral, highly selective, brain penetrant, small molecule inhibitor of Elovl1 - CPD37.

CPD37 had favorable pharmacological properties, with strong potency on target (IC50 = ∼50 nM), sufficient oral bioavailability of 46% and a brain to plasma ratio of 1.1. With daily oral doses of 100 mg/kg in mice, it achieved ∼20 hours of coverage over the plasma free fraction corrected IC_50_. A high level of inhibition was necessary for VLCFA reduction, as only the 100 mg/kg daily dose was able to achieve full normalization of C26:0 in the plasma in as little as 7-days of dosing and sustained over a 30-day dosing period. Yet, little if any changes in C26:0 was apparent in the CNS after 30-days of dosing. Indeed, this amount of compound given in feed was also able to achieve substantial reductions of C26:0 in tissues after 6 months. After prolonged treatment, liver and kidney had WT-like levels of VLCFAs, while the brain and spinal cord had 70% and 66% reduction respectively. These changes were unlikely to be driven by the halting of further accumulation of VLCFA in the CNS, since treatment was started at 6 months of age. In our characterization study, brain C26:0 levels were largely stable after 6 months. While there was continuous accumulation of C26:0 in the spinal cord throughout the lifetime of the animal, the bulk of the accumulation occurred before 6 months. We expect that complete normalization of VLCFAs may be possible with longer treatment.

While the goal of VLCFA normalization through Elovl1 inhibition seems possible, would this therapeutic approach correct the pathology associated with loss of *ABCD1*? After all, studies with alternative therapeutic agents such as dimethyl fumarate have demonstrated a benefit in mouse models without affecting VLCFA levels.^37^ In the *ABCD1^−/y^* mice many of the documented behavioral phenotypes are only apparent in very aged animals and our behavioral analyses confirmed the absence of consistent behavioral deficits in the first year of the animal’s life.^31^ Instead, we turned to RNA sequencing to identify early cell-type dependent transcriptional changes, which may additionally be useful biomarkers for successful therapeutic intervention. We performed single nuclei sequencing of *ABCD1*^−/y^ mice, at 12 months, with and without Elovl1 inhibitor treatment. While there were some downregulated pathways in the brain, mostly in astrocytes, oligodendrocytes, and endothelial cells, here too, the differences between WT and *ABCD1*^−/y^ were subtle. Changes in the spinal cord were even less pronounced and largely restricted to just oligodendrocytes, with some involvement of dopaminergic neurons and endothelial cells. To our surprise, treatment with Elovl1 inhibitor led to much more profound transcriptional changes in genes beyond those altered by loss of Abcd1. Most prominently, broad pathway perturbations were observed in cholinergic and glutaminergic neurons in the spinal cord, including increased activity of protein unfolding stress, ERAD signaling, and translation. These same pathways were otherwise unaffected by *ABCD1*^−/y^. Taken together, these results suggest that CPD37 treatment, and more broadly Elovl1 inhibition, may have unexpected consequences in *ABCD1*^−/y^ mice that go beyond correction of lipid homeostasis.

Brain-penetrant Elovl1 inhibitors with chemical structures highly distinct from ours have been reported previously.^25,26^ The PK/PD relationship of these other compounds mirrored that of our own. VLCFA levels in the blood normalized within a week of daily treatment, followed by a longer-term decrease in the CNS with 66% reduction after 8 months. While those studies measured only Lysophosphotidylcholine C26:0 levels, our measurements of fatty acid levels were more complete. By hydrolyzing the acyl chain of all lipids, we reported decreases in total C26:0 fatty acids across all lipid species. These data support the hypothesis that Elovl1 inhibition can effectively reduce very long chain fatty acids in the CNS, across a broad spectrum of lipid species. Reduction of VLCFAs in the CNS required long-term treatment with CPD37 at high doses, suggesting that strong inhibition needs to be maintained for a long period of time to achieve the goal of normalizing VLCFAs in the organs relevant for ALD. This is likely due to the slow turnover rates of sphingolipids^38^, which are a major component of CNS lipids.^39^

Despite efficacious target engagement, the therapeutic index of our compounds and those reported in the literature were insufficient to warrant clinical investigation. Yet, there are differences in the target organs for both sets of compounds, which may point to the possibility that the findings are off target or species specific. For both the pyrimidine ether-based compound 22 and pyrazole amide-based compound 27, corneal opacities were a shared finding in adult rats.^25,26^ Interestingly, in our preliminary toxicology studies in rodents, there were no adverse effects in the eye. For CPD37 the main dose limiting toxicities were related to the digestive tract and the skin. These digestive tract findings may be a common link between our compound and previously reported compounds, however for all three compounds it was most prominent in the forestomach, which is unique to rodents. In NHPs, stomach and ocular findings with compounds 22 and 27 were not observed.^25,26^ Instead, the major findings for compound 22 were in the eyelid and skin areas not covered by hair, combined with convulsions at high doses. The skin findings are perhaps the most concerning for applications in humans. ELOVL1 knockout mice have defects in skin barrier function, which is neonatal lethal.^27^ Relatedly, in a handful of human case studies with *ELOVL1* mutations, patients presented with ichthyosis and acanthosis nigricans, consistent with a loss of moisture retention.^40–42^ Yet, our analysis of the heterozygous *ELOVL1* knockout mouse revealed no findings in these organs and many people carrying likely loss-of-function mutations in *ELOVL1* can be found in the gnomAD database of humans with no apparent illnesses. These data suggest Elovl1 may indeed be a tractable target for therapeutic intervention, but the data reported here and with other Elovl1 inhibitors suggest it is a challenging one.

The long duration of Elovl1 inhibition required to reduce VLCFAs in the *ABCD1*^−/y^ animals raises the question of whether there is insufficient degradation capacity in this genotype or if the lipids are in immobile pools. The small changes in VCLFA following the treatment of WT animals with Elovl1 inhibitors for 12 days supports the latter hypothesis. Indeed, when we performed lipidomic analyses following 30 days of Elovl1 inhibition in *ABCD1*^−/y^ animals, there was a dramatic difference in the reduction of some lipid classes over others. Namely, phosphatidylcholines as a class were significantly reduced after 30-days of treatment, while sphingolipids and glycosphingolipids were hardly changed. Stable isotope labeling-based measurements of mouse myelin lipid turnover produced similar observations of the relatively quicker turnover of phosphatidylcholines.^43^ The sensitivity of phosphatidylcholines to perturbations in the synthesis or degradation machinery of lipids make them an ideal candidate for target engagement biomarkers for molecules like Elovl1 inhibitors. Moreover, the differences in the relative rates of lipid changes following Elovl1 inhibition may point to a path forward for Elovl1 inhibitors. If changes in lipid species which are more sensitive to Elovl1 inhibition might be disease modifying for ALD, then an acceptable TI might exist. The potential benefit of Elovl1 inhibition may extend beyond ALD as well, considering Elovl1 substrates have been postulated as mediators of neuroinflammation.^44^ Identifying a particular lipid or groups of lipids that underlie the pathology of these diseases, if there are such specific lipids, could be greatly enabling for the development of novel metabolism modulating medicines.

## Acknowledgements

The authors would like to thank Dr. Aurora Pujol for her guidance on the project, Andre Kurlovs for his assistance in data management, Dr. Marc Engelen and Dr. Troy Lund for their consultations.

## Author contributions

JYH analyzed and interpreted the single nuclei sequencing data, prepared figures, wrote and edited the manuscript. BF led chemistry efforts, conceptualized and supervised compound profiling and edited the manuscript. DSB, MSRC, AB, YL, KK, ZZ, SNK, ST, HM, JJ, designed, executed, and analyzed *in vivo* experiments. AM led drug metabolism and pharmacokinetics efforts. AB and JD led preclinical safety studies. YL led the measurement of drug concentration in animal studies. CP performed the CETSA experiments, analyzed, and interpreted the data. JDP conceptualized, executed, analyzed the long-term lipid profiling of the mice, and edited the manuscript. DF contributed to the design and synthesis of the chemical matter. CSS led the scale-up synthesis for pharmacology and preclinical safety studies. SV developed formulations for *in vivo* studies. AG conceptualized, designed and executed elongase assays; analyzed the data. LP and LG designed and executed lipidomics experiments, analyzed data and prepared figures. BZ supervised lipidomics and CETSA work. AB and KR developed lipid detection analytical methods for cell-based compound profiling. RM was the project manager. MH, DW, designed and executed *in vitro* compound profiling, analyzed data and prepared figures. AK conceptualized and supervised global lipidomics and fatty acids profiling and edited the manuscript. BL, SH, ZD, BT designed, executed in vitro compound profiling experiments, analyzed data, and prepared figures. TG led multidisciplinary project team, conceptualized and supervised in vitro compounds profiling and edited the manuscript. JG processed tissues, generated single nuclei sequencing data, and analyzed the data. ET analyzed single nuclei sequencing data. GG analyzed single nuclei sequencing data and supervised the analysis team. SPS supervised the project and the teams. CK led biology efforts, conceptualized, executed, and supervised in vivo experiments, analyzed data, produced figures, and wrote the paper.

---

**Figure S1.**
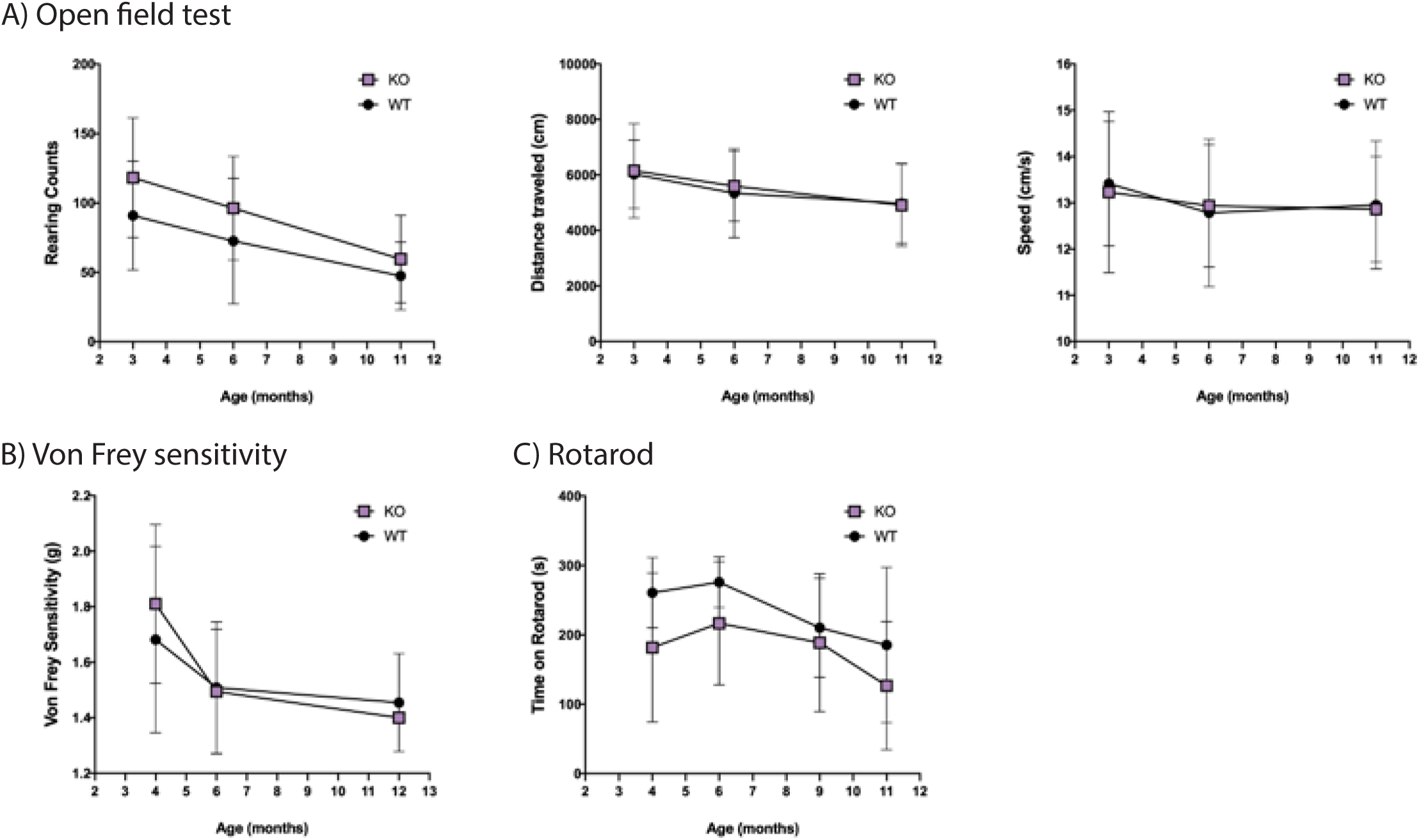
Behavioral analyses including **A)** open field test, **B)** von Frey sensitivity, and **C)** rotarod as a function of age. N = 20-23 male animals. No differences were statistically significant by two-way ANOVA.

**Figure S2.**
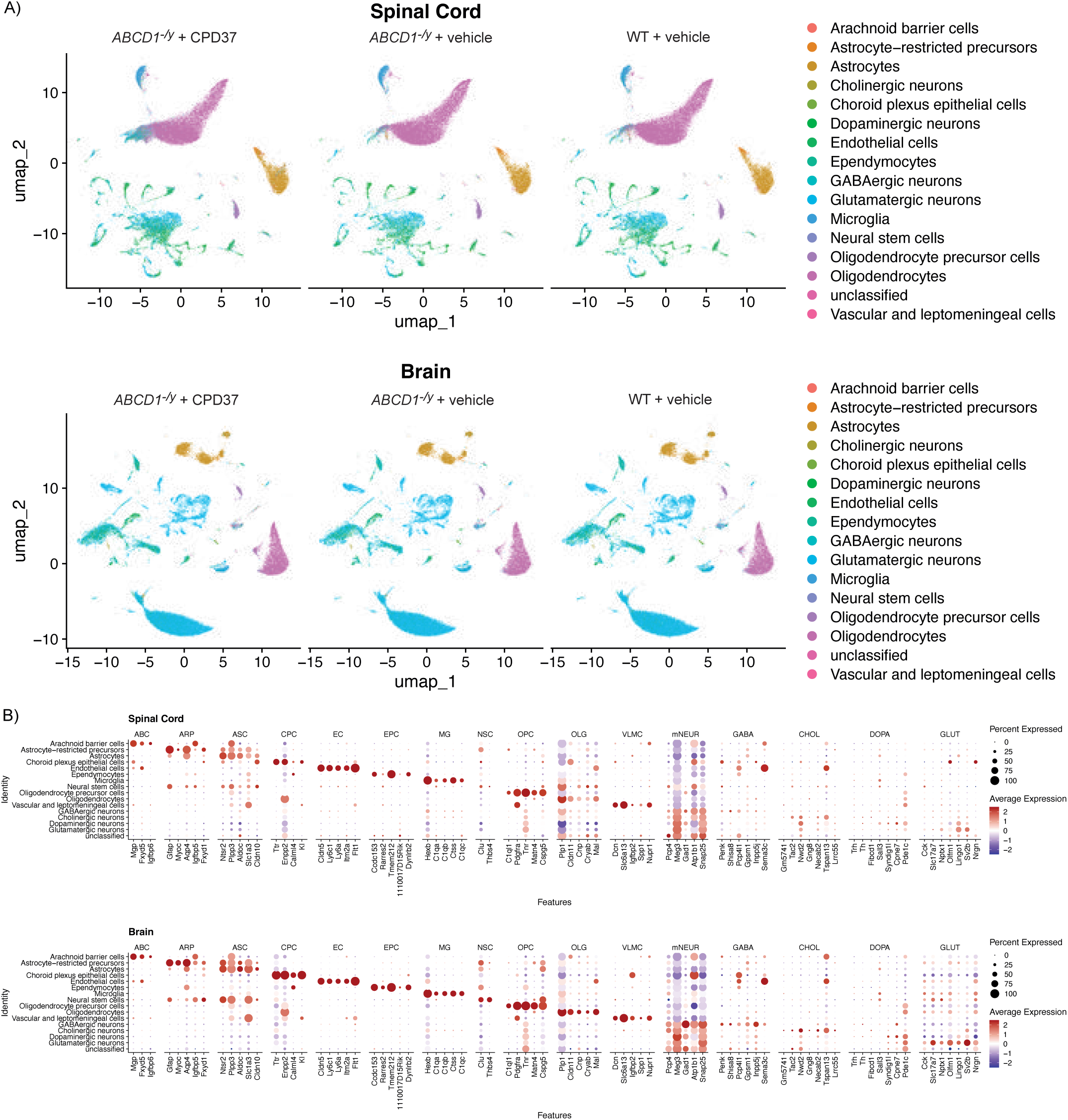
Clustering and Cell Type Annotation of Wild-Type, *ABCD1^−/y^*, and *ABCD1^−/y^* + Treatment Mice: **A)** UMAP dimensional reduction plot comparing the cells between different mouse models in the (top) spinal cord and (bottom) brain. **B)** DotPlot with ‘top 5’ markers for each of the cell types that were used to perform cell type calling via the SARGENT algorithm in the (top) spinal cord and (bottom) brain.

**Figure S3:**
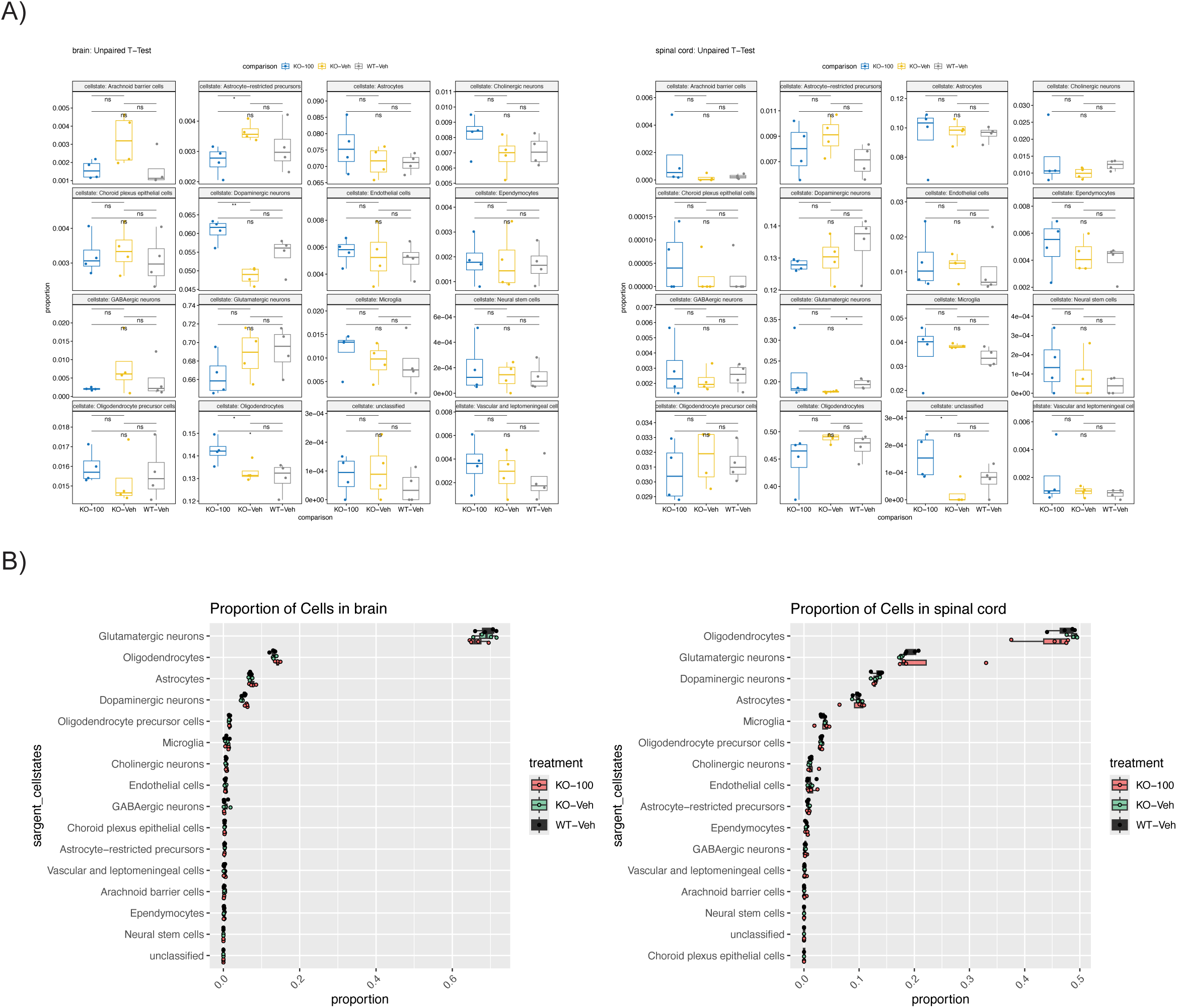
Comparison of Cell Type Proportions in Wild-Type, *ABCD1^−/y^*, and *ABCD1^−/y^* + Treatment Mice: **A)** Boxplots comparing the relative proportion of each cell type across treatment groups in (left) brain and (right) spinal cord. Unpaired t-test: * p < 0.05, ** p<0.005. **B)** Boxplots referencing total proportion of cell types in each sample across (left) brain and (right) spinal cord.

**Figure S4:**
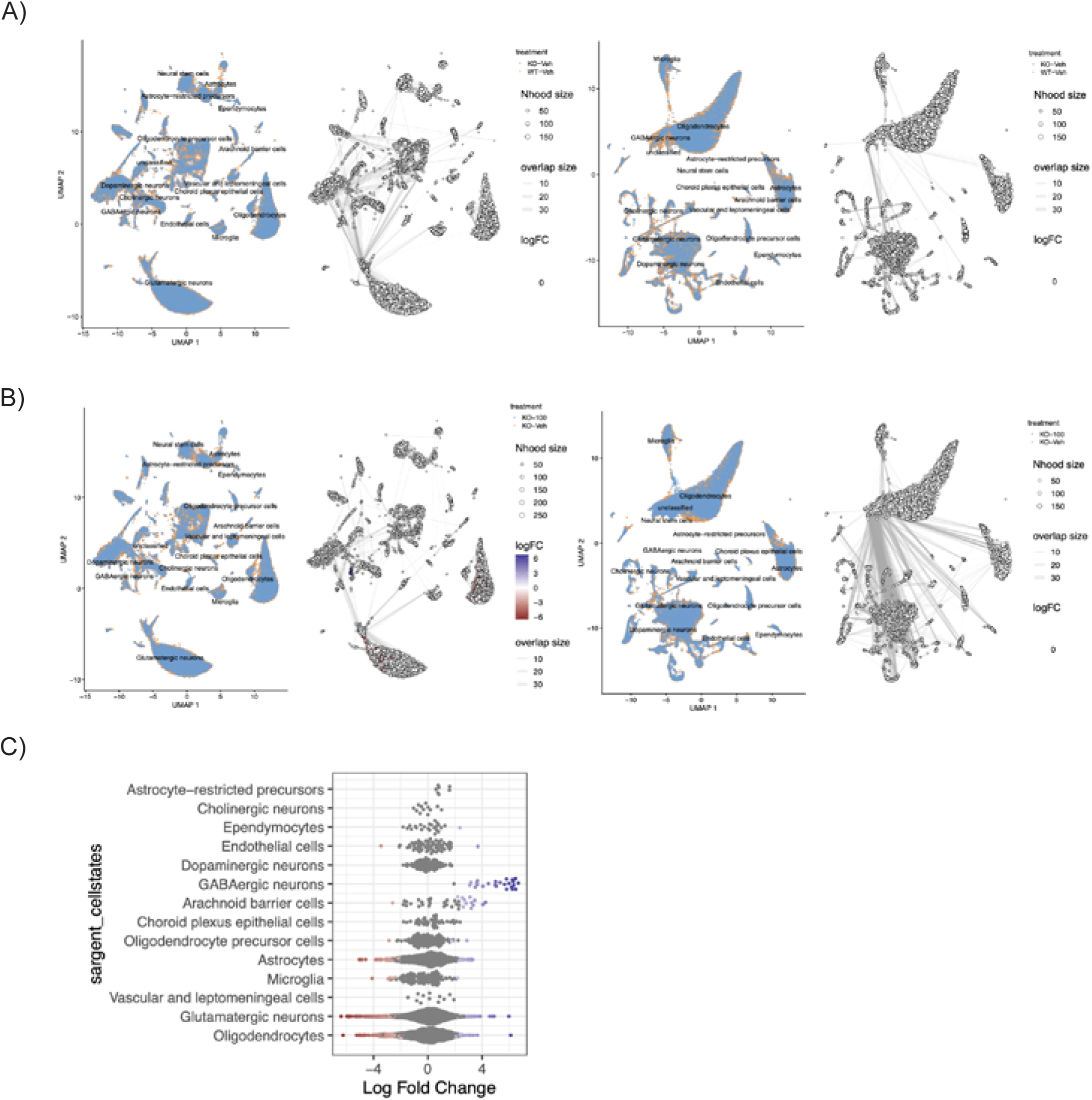
Comparison of Differentially Abundant Cellular Neighborhoods by KNN in Wild-Type, *ABCD1^−/y^*, and *ABCD1^−/y^* + Treatment Mice: **A)** UMAP visualizing cellular neighborhoods in Abcd1-/^y^ and Wild Type group comparison in (left) brain vs (right) spinal cord. No statistically significant enrichment detected in differential abundance with a Spatial FDR cutoff of 0.25. **B)** UMAP visualizing cellular neighborhoods in CPD37 + *ABCD1^−/y^* and *ABCD1^−/y^* group comparison in (left) brain vs (right) spinal cord. Cellular neighborhoods which are differentially abundant in the CPD37 treatment group are colored in blue versus those enriched in *ABCD1^−/y^* only group are highlighted in red. **C)** Beeswarmplot showing the SARGENT labeled identities of enriched cellular neighborhoods in the brain.

**Figure S5:**
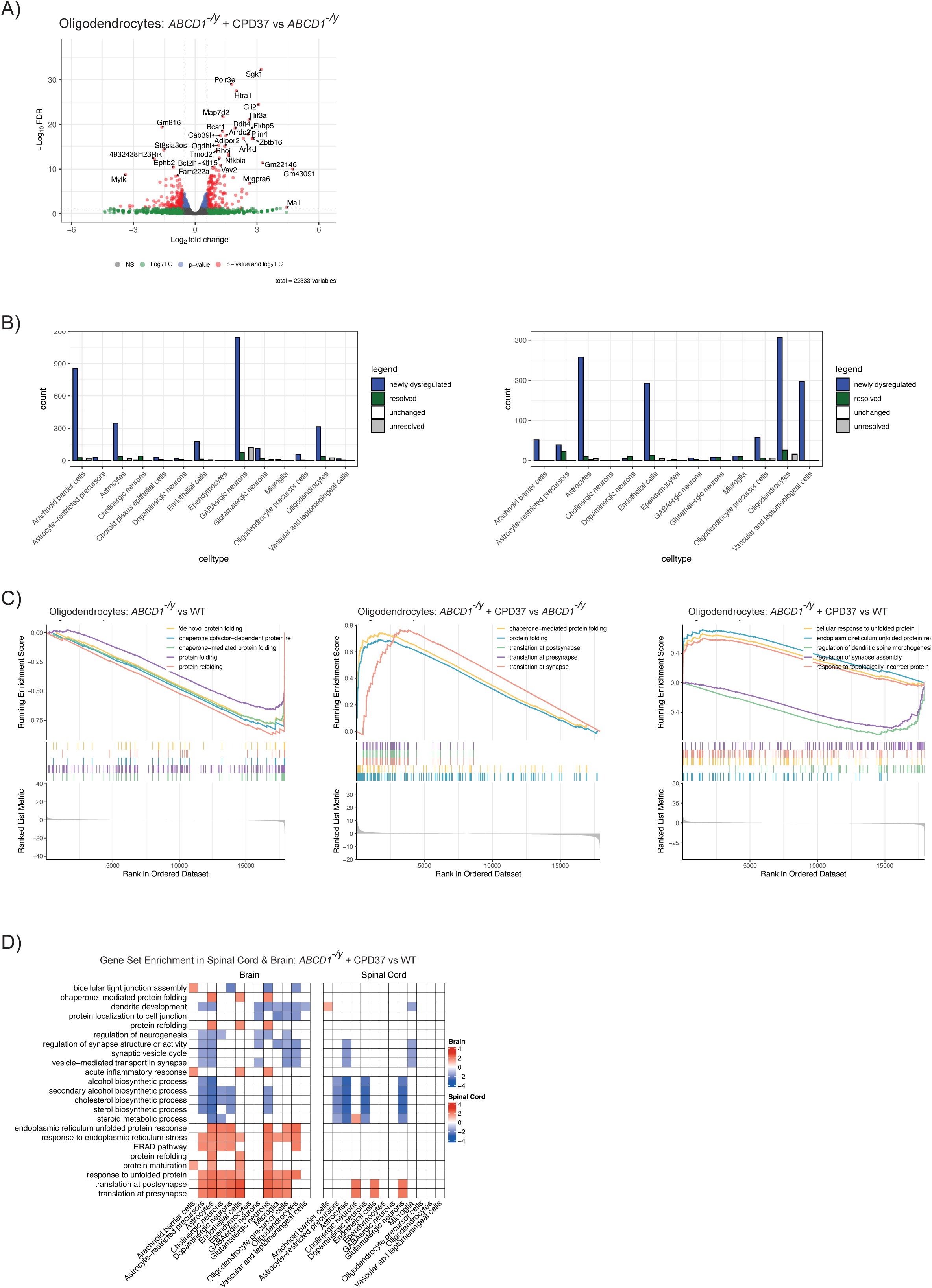
Comparison of Differentially Expressed Genes in Wild-Type, *ABCD1^−/y^*, and *ABCD1^−/y^* + Treatment Mice: **A)** Volcano plot of the differential expression of genes in a comparison of *ABCD1^−/y^* mice compared to Wild-Type in Oligodendrocytes in the brain. **B)** Paired barplots showing the number of genes which are newly dysregulated by CPD37 treatment in *ABCD1^−/y^* mice versus resolved towards wildtype, unchanged, or unresolved in (left) brain or (right) spinal cord. **C)** Gseaplot2 plots denoting the top 5 enriched gene ontology biological process genesets enriched in geneset enrichment analysis in the brain in (left) *ABCD1^−/y^* vs WT, (middle) CPD37 + *ABCD1^−/y^* vs *ABCD1^−/y^*, and CPD37 + *ABCD1^−/y^* vs WT comparisons. **D)** Heatmap which shows enriched pathways in a comparison of CPD37 + *ABCD1^−/y^* mice vs Wild Type vehicle controls across cell types.

## Methods

### Generation of Elovl1 heterozygous knockout mice

We used CRISPR-Cas9 gene editing to generate the *ELOVL1*^−/+^ mouse with the deletion of the entire coding region of one copy of *ELOVL1*. The gene targeting was designed and performed by Jackson Laboratories (Bar Harbor, ME). Briefly, two sgRNAs (Elovl1_intron: ACATCGGCTGCCGAAGCATC, Elovl1_UTR: TCTACCTACACCCGTGACCA) and guide RNA were introduced into C57Bl/6J embryos. 5 Founder animals (F0) were screened for the deletion by PCR amplification of tail DNA using two sets of primer pairs: Elovl1_genoF1: TTTACCTGGGAGAAGGGGGT, Elovl1_genoF2: TCTTGGAAAAGGGCCAGAGG, Elovl1_genoR1: TTCAGCCTCCAGTTCTGGTG, Elovl1_genoR2: AGGAGAAAGGCCACAAGACC. The founder with a 1958 base pair deletion was selected for breeding and crossed to C57Bl/6j to produce study cohorts.

### Animals

The Institutional Animal Care and Use Committee (IACUC) at Sanofi approved all procedures. All experiments were performed in accordance with relevant guidelines and regulations as stipulated by the IACUC. Animals were housed under light:dark (12:12 h) cycles and provided with food and water ad libitum. Environmental enrichment was provided in the form of corn cob bedding, enviro-dry, nestlets, and cardboard nest boxes. Dietary enrichment was not provided to any groups. All behavioral testing was performed during the animals’ light cycle, between the hours of 8 AM and 4 PM. All methods and data are reported in accordance with ARRIVE guidelines (https://arriveguidelines.org) and recommendations.

### Administration of Elovl1 inhibitor (CPD37) and control

For all treatment studies that were 30 days or shorter, the compound was formulated in a nanosuspension (0.5% Methylcellulose 0.2% Tween 80 in water or 2%PVP k25, 0.1%SLS in water) and was administered via oral gavage at 5 ml/kg. For the 6-month treatment study, CPD37 was compounded into rodent chow (Research diets, catalog # D11112201) at a concentration of 0.05% (w/w) for an approximate daily dose of 100 mg/kg. Control mice were fed matched chow without compound. CPD37 and control diets were provided *ad libitum* until necropsy.

### Animal perfusion and tissue and blood collection

Mice were anesthetized with isoflurane and ∼500 mL of whole blood was collected from the retro-orbital sinus using a glass capillary tube into a Microtainer® tube (BD Biosciences; Billerica, MA) containing K_2_ EDTA anticoagulant. Plasma was isolated after 10 min centrifugation at 3200 RPM at 4 °C. Immediately following blood collection, animals were euthanized by C02 asphyxiation. Where needed, following euthanasia mice were transcardially perfused with cold phosphate-buffered saline (PBS) at a rate of 18 ml/minute, for two minutes. After dissecting the brains sagittally along the midline, the left hemisphere was further dissected into regions. For the characterization study (Figure 5), the cortex was isolated. For all other studies, the rostral half of the hemibrain brain was used. Whole spinal cord and 1 whole kidney were used for the relevant analyses, Figure X also included a piece of a single lobe of the liver. All tissues were weighed and frozen in Lysing Matrix D tubes (MP Biomedicals #6913100) on dry ice and stored at − 80 °C until use.

#### Histology and Histopathology

Representative samples tissues from 6-month-old Elovl1 KO heterozygote mice and WT control mice were collected and fixed in 10% neutral buffered formalin. Formalin-fixed tissues were processed for paraffin embedding and sectioned at 5 micron and stained with hematoxylin and eosin (H&E). H&E slides were scanned into digital whole slide images (WSIs) using Leica GT450 scanner (Leica Biosystems). WSIs were examined by a veterinary pathologist blinded to the study groups.

### Measurement of Very Long Chain Fatty Acids (VLCFA) levels

Brain and spinal cord tissue was homogenized in water for a final concentration of 200mg/ml. Fatty acid profiling was performed for total fatty acids including free fatty acids and bound fatty acids after derivatization with methanol in presence of Boron Trifluoride (BF3) as a catalyst. The derivatization products, Fatty Acid Methyl Ester (FAME) were analyzed by GC-MS/MS. Total 35 fatty acids were included in this method including saturated, mono-unsaturated, and poly-unsaturated fatty acids. The solid tissue samples (spinal cord, stomach, brain, and skin) were then homogenized in deionized water for a final concentration of 200 mg/mL. All samples were stored at -80°C until analysis.

For the analysis of plasma samples, 50 µL of plasma sample was transferred into a digestion tube, and added 2 mL BF3 in Methanol and 40 µL of internal standard solution (250 µg/mL of C19:0 methyl ester with 1200 µg/mL BHT in heptane). The samples were then digested on a heatblock at 100°C for 1 hour. Once cooled, 2 mL heptane and 4 mL DI water were added for extraction. An aliquot of the heptane phase was transferred into an autosampler vial for analysis by GC-MS/MS using GC: Agilent 7890A GC coupled with Agilent 7000C MS/MS).

For the analysis of tissue samples, 250 µL of sample homogenates was transferred into a digestion tube, and 2 mL BF3 in Methanol and 40 µL of internal standard solution (250 µg/mL of C19:0 methyl ester with 1200 µg/mL BHT in heptane) were added. The samples were then digested on a heatblock at 100°C for 1 hour. Once cooled, 2 mL heptane and 4 mL DI water were added for extraction. Sample extracts were analyzed undiluted and diluted 1:50 with diluent by GC-MS/MS. Calibration standards for all 35 fatty acid methyl esters (FAMEs) were prepared in heptane with the internal standard (C19:0 methyl ester).The analysis method employed was capillary gas chromatography with chemical ionization (CI) and detection with a triple-quadrupole mass spectrometer (GC-MS /MS ). Methane gas was used for the chemical ionization.

Fatty acids were quantified using calibration curves that consisted of at least 5 concentration levels. Calibration curves were established by a regression of either linear or quadratic fit with or without 1/x weighing and required for a minimum R2 greater than 0.99. The back-calculated calibration standards were within 25% of the nominal values. Data was processed using MassHunter 8.0 software.

### Rotarod test

Motor coordination and balance testing was performed using the UGO Basile Rota-Rod. The first timepoint encompasses additional days to acclimate and condition the animals. Each testing day comprised of three trials with at least one hour rest in between trials. On day one “Acclimation”, the rod spun at a constant speed of 5RPM over 120seconds with replacement if the animal fell. One day 2 “Conditioning”, the rod accelerating from 5-20RPM in either direction over 300 second without replacement. Day three was testing day where average fall latency from the rod (or cutoff time) was recorded for each animal using the same settings as day 2. For subsequent timepoints, only the day 3, the testing day was performed.

### Von Frey sensitivity test

Mice were placed in enclosed compartments on a raised grid and allowed to acclimate for ∼10 minutes. Next, the von Frey fibers were applied to the middle plantar surface of the hindpaw starting at a 1.4g fiber force, followed by the 2.0g fiber force.

If the mouse withdrew or licked the paw soon afterwards, it was considered a positive sensation of the fiber. A total of 10 attempts were made with a given fiber strength alternating between the paws. The force required to induce a withdrawal in 50% or more of the attempts was reported as the sensitivity value. If fewer than 50% of the attempts were sensed, the next fiber strength was tested until the individual reached a 50% sensitivity threshold.

### Photobeam Activity System (PAS) Open Field locomotion assay

San Diego Instruments PAS-Open Field stations use a 16 x 16 photobeam configuration to measure spontaneous and exploratory behavior in mice. On testing days, mice are acclimated to the behavior suite for 30 minutes prior to testing. Mice are placed in individual plexi glass chambers and allowed to freely explore. Total distance, rearing and speed is measured over 20 minutes sessions.

### Stable isotope labeled elongation assay

Stable isotope labeled stearic acid (^13^C-18) was purchased from Sigma (Cat# 605581) and formulated in 100% hemp oil at a concentration of 50 mg/ml. C57Bl/6 (Charles River Laboratories) M/F mice, ages P17-20 were dosed PO 10 mL/kg BW with CPD37, followed by a 100 uL dose of the 50 mg/ml ^13^C-18 one hour later. This process was repeated for three days. On the third day of dosing, the mice were sacrificed 4 hours after the dose of ^13^C-18. Tissue samples were collected and fatty acid analysis was performed by GC-MS as described above.

### Untargeted lipidomics

50 µL tissue homogenates (200 mg/mL in water) and blanks were transferred to a microcentrifuge tube containing 900 µL 1:1 acetonitrile:methanol spiked with internal standards. Samples were vortexed at room temperature for 15 min, then centrifuged for 6 min at 10,000 rpm. 900 µL of supernatant was transferred to an HPLC vial and placed under a stream of nitrogen gas until the solvent was evaporated. Dried samples were reconstituted in 90 µL of 7:3 acetonitrile:methanol with brief vortexing. Samples were then transferred to QSert HPLC vials, with 10 uL reserved to create pooled QC samples for the separate tissues. 2 µL of sample were injected for LC-MS analysis using a Waters Acquity UPLC system coupled to a Thermo Scientific Q-Exactive mass spectrometer. LC was performed at a constant flow rate of 0.3 mL/min on a 2.1 x 150 mm, 1.7 µm Waters Acquity BEH HILIC column held at 30 °C. The method started with 2 min at 95% Mobile phase A (96% acetonitrile, 2% methanol, 1% water, 1% acetic acid, 5 mM ammonium acetate) and 5% Mobile phase B (98% methanol, 1% water, 1% acetic acid, 5 mM ammonium acetate) followed by a gradient over 13 min to 50% B, followed by a gradient over 3 minutes to 40% B. The system was held at 40% B for 1 min then switched to 5% B and held for 6 min. Full MS data was acquired in a single injection per sample using positive/negative polarity switching. Raw data were processed using Genedata Expressionist v. 17.5.4 Refiner MS. Data were filtered to exclude features with relative standard deviation in pooled QC sample >30%. Statistical analysis was performed using Metaboanalyst v.6.0. Variables with >70% missing values were removed, and the remaining missing values were imputed as 1/5th the minimum of the variable value. Log transformation and scaling (mean-centered and divided by the standard deviation of each variable) was applied to the data which were analyzed by t-tests.

### Single-nuclei RNA sequencing data

*Nuclei Isolation:* Tissue samples were stored at -80°C. For tissue lysis and washing of nuclei, sample sections were added to 1 mL Nuclei PURE lysis buffer (Sigma, NUC-201) and thawed on ice. Samples were then Dounce homogenized with PestleAx20 and PestleBx20 before transfer to a new tube, with the addition of additional lysis buffer. Following incubation on ice for 15 minutes, samples were then filtered using a 30 mM MACS strainer (Fisher, NC0642012), centrifuged at 500xg for 5 minutes at 4°C using a swinging bucket rotor (Sorvall Legend RT, Thermo Fisher), and then pellets were washed with an additional 1 mL cold lysis buffer and incubated on ice for an additional 5 minutes.

A 1.8M sucrose buffer was created and 1.8ml was mixed with 1ml of cell suspension. 1ml of sucrose buffer was deposited to ultra centrifuge tube (Fisher, NC9157569) then the cell suspension was layered on top (2.8ml). The suspension was then centrifuged at 30,000xg (17,800 rpm) for 45min at 4C, Rotor SW55ti. Supernatant was then aspirated off, and pellet was resuspended in 1ml of wash buffer (Nuclei Pure storage buffer, Sigma NUC-201) and spun at 500xg for 5 min at 4°C. Supernatant was aspirated and 0.5ml of wash buffer was added for resuspension.

For NeuN/Dapi and FACS sorting, from 0.6 mL nuclei sample, 540 mL, 30 mL, and 30 mL were aliquoted into tubes for sample and controls and then 10X Dapi/NeuN buffer was added to tubes for a final 1X concentration. Tubes were then incubated on ice for 30 minutes, with inversion every 10 min. Following incubation, samples were spun at 500xg for 5 min, supernatant removed, and samples were resuspended in 600 ul Wash buffer for samples (300 ul for control tubes). Nuclei then underwent filtering through 70um filter and filtered again through 40um filter (Fisher, 14100150) just before sorting using BD Bioscience InFlux Cell Sorter.

#### Sample QC and library Preparations

Library preparation and NovaSeq Sequencing: Libraries were prepared according to 10xGenomics protocol for Chromium Single Cell 3’ Gene Expression V3.1 kit. NovaSeq sequencing was performed according to illumine NovaSeq 6000 protocol. UMI count matrices generated by Cellranger V5.0.0.

#### Single-nuclei RNA sequencing data preprocessing

CellBridge^32^, was used to perform pre-processing from HDF5 format to a packaged Seurat object for the OTAR2057 dataset. CellBridge, in brief, translates and packages HDF5 files into R-based Seurat object and performs QC thresholding (minimum UMI = 750, percent.mt < 20, nFeature_RNA<250), a log normalization of count data (LogNormalize function with a scaling factor of 1e+06), FindVariableFeatures (nfeatures=2000), RunUMAP (dims = 1:30, res=0.7, k=20), and Harmony (by sample). Automated celltype calling was performed by SARGENT, a cell-by-cell geneset-based classification algorithm. Genesets for cell-type annotation were acquired from previously annotated cell types^34^ from supplementary table 3 (cell-type marker genes) and 4(neuronal subtype marker genes) and filtered for the top positive (log2FC > 1.5, pct.1/pct.2 > 10 | pct.1-pct.2 > 0.5, slice_max(n=50) and negative markers (log2FC < - 1.5, pct.2/pct.1 > 10 | pct.2-pct.1 > 0.5). Discrimination of mature neural subtypes (Gabaergic, Cholinergic, Dopaminergic, and Glutamatergic) were annotated by a second application of SARGENT using the following criteria (log2FC > 1.5, pct.1/pct.2 > 2.5 | pct.1-pct.2 > 0.5, slice_max(n=25)). All code for cell type calling and processing to be provided in Sanofi-Public github.

#### Analysis Pipeline

Primary data visualization pipelines in R were run under the R 4.3.2 environment. UMAPs were plotted using native Seurat (5.1.0) functions. Cell type proportion comparisons were visualized using the ggboxplot function in the ggpubr (0.6.0) package and significance comparisons were performed using an unpaired t-test for the cell types considered in the plot. For differential expression analyses between conditions, a custom wrapper function using edgeR (4.0.16)/limma (3.58.1) was created. In brief, this function performs pseudobulk aggregation using the function AggregateExpression on a Seurat object’s raw count data, per condition_cellstate. Then, we created a pairwise design and contrast matrix and the equation ∼0+comparsioncellstates with no input covariates into the analyses. A standard recommended pipeline for creating a negative binomial generalized linear model with F-test using filterByExpr, calNormFactors, estimateDisp, and glmQLFit was then called and individual glmQLFTests were run in parallel using foreach (1.5.2) and doParallel (1.0.17) packages. Differential expression results were visualized using the EnhancedVolcano (1.14.0) package and Gene Set Enrichment Analysis was performed on a targeted list of genesets using a custom wrapper function which uses bioMart (2.52.0) and org.Mm.eg.db, GSEA function and Gene Ontology: Biological Process using the gseGO functions within the ClusterProfiler (4.10.0) package. Gseaplot2 was used to visualize the enrichment plots and rank was calculated by sign(logFC) * -log10(PValue) based on edgeR standard outputs. For differential abundance analyses between conditions, we used the miloR (1.4.0) package following the recommended vignette tutorial with an FDR (alpha) cutoff of 0.25. Beeswarm Plots were generated via plotDAbeeswarm function, UMAPs of key neighborhoods were plotted using plotNhoodGroups,

##### 14-day repeat dose rat toxicology study

In vivo toxicology studies were conducted by Sanofi, Montpellier France. CPD37 was administered once daily to Crl:CD (SD) rats by oral gavage for up to 14 days at dose levels of 0, 50, 150, or 450 mg/kg/day at a dose volume of 10 ml/kg. Compound was formulated as a microsuspension in 0.5% MC/0.2% Tween 80 in water. Male rats were obtained from Charles River Lab, Italy and were approximately 6-7 weeks of age on Day 1 of dosing. Five male animals were assigned to each group. The following endpoints were evaluated: clinical signs, body weight, food consumption, TK (serum, liver), clinical pathology (hematology, clinical chemistry, coagulation, urinalysis), histopathology, and biomarkers.

##### Human Elongase enzymes expression

Elovl 1, 3, 6 and 7 were expressed using Baculovirus system; full lengths ELOVLs proteins were cloned in pFastBac3 (or 5) vector with N-terminus His-Flag tags (BPS Biosciences).

ELOVL1 (aa 2-279(end)) NM_022821:

MHHHHHHGSSGDYKDDDDKGSSGENLYFQGSSGEAVVNLYQEVMKHADPRIQGYPLMGSPLLMTSILLTYVYFVLSLGPRIMANRKPFQLRGFMIVYNFSLVALSLYIVYEFLMSGWLSTYTWRCDPVDYSNSPEALRMVRVAWLFLFSKFIELMDTVIFILRKKDGQVTFLHVFHHSVLPWSWWWGVKIAPGGMGSFHAMINSSVHVIMYLYYGLSAFGPVAQPYLWWKKHMTAIQLIQFVLVSLHISQYYFMSSCNYQYPVIIHLIWMYGTIFFMLFSNFWYHSYTKGKRLPRALQQNGAPGIAKVKAN

ELOVL3 (aa 2-270(end)) NM_152310:

MHHHHHHGSSGDYKDDDDKGSSGENLYFQGSSGVTAMNVSHEVNQLFQPYNFELSKDMRPFFEEYWATSFPIALIYLVLIAVGQNYMKERKGFNLQGPLILWSFCLAIFSILGAVRMWGIMGTVLLTGGLKQTVCFINFIDNSTVKFWSWVFLLSKVIELGDTAFIILRKRPLIFIHWYHHSTVLVYTSFGYKNKVPAGGWFVTMNFGVHAIMYTYYTLKAANVKPPKMLPMLITSLQILQMFVGAIVSILTYIWRQDQGCHTTMEHLFWSFILYMTYFILFAHFFCQTYIRPKVKAKTKSQ

ELOVL6 (aa 2-265(end)) NM_024090:

MHHHHHHGSSGDYKDDDDKGSSGENLYFQGSSGNMSVLTLQEYEFEKQFNENEAIQWMQENWKKSFLFSALYAAFIFGGRHLMNKRAKFELRKPLVLWSLTLAVFSIFGALRTGAYMVYILMTKGLKQSVCDQGFYNGPVSKFWAYAFVLSKAPELGDTIFIILRKQKLIFLHWYHHITVLLYSWYSYKDMVAGGGWFMTMNYGVHAVMYSYYALRAAGFRVSRKFAMFITLSQITQMLMGCVVNYLVFCWMQHDQCHSHFQNIFWSSLMYLSYLVLFCHFFFEAYIGKMRKTTKAE

ELOVL7(aa 2-281(end)) NM_024930:

MHHHHHHGSSGDYKDDDDKGSSGENLYFQGSSGAFSDLTSRTVHLYDNWIKDADPRVEDWLLMSSPLPQTILLGFYVYFVTSLGPKLMENRKPFELKKAMITYNFFIVLFSVYMCYEFVMSGWGIGYSFRCDIVDYSRSPTALRMARTCWLYYFSKFIELLDTIFFVLRKKNSQVTFLHVFHHTIMPWTWWFGVKFAAGGLGTFHALLNTAVHVVMYSYYGLSALGPAYQKYLWWKKYLTSLQLVQFVIVAIHISQFFFMEDCKYQFPVFACIIMSYSFMFLLLFLHFWYRAYTKGQRLPKTVKNGTCKNKDN

All ELOVL proteins were expressed in SF9 insect cells by infection of 1 ml of baculovirus (titer 3-5*10^8^-1*10^9^ pfu/ml) to 1 L of cultured media (Expression Systems, ESF921) and the cells were harvested 72 h later with cell viability >75-80%. Average cell pellet weight was 7-15 g per 1 L of SF9 cells culture. Cells were washed with PBS buffer (Invitrogen, 20012) and stored at -80^0^C in 50 ml conical Falcon tubes.

##### ELOVL microsomes preparation

During the preparation of microsomes all steps were carried out in a chilled environment.

Pellet of insect cells expressing the ELOVL proteins (pellet size ranging from 15-35g/2L culture) kept in 50ml Falcon tube at -80^0^C was thawed in a cold water bath at 4^0^C while spinning; and added to 200 ml of prechilled microsome buffer (50mM Hepes pH 7.5 (Sigma H0887), 25mM KCl (Sigma P3911), 0.25M sucrose (Sigma S0389), 1 tablet per 50ml protease inhibitor (Roche, COUEDTAF-RO), 0.1mM sodium orthovanadate (Sigma, S6508, 0.1mM beta-glycerol phosphate (Sigma, G6251), and 20% glycerol (Sigma, G2025) when the pellet was completely thawed. Cells suspension was next sonicated on ice using a probe sonicator (Branson) with a horn tip. Suspension was spin down at 10,000 g (Beckman, J-30I) for 10 minutes at 4^0^C in 6 pre-chilled Nalgene round bottom tubes followed by supernatant spin down at 100,000 g (Beckman, J-30I) for 1 hour at 4^0^C in Beckman round bottom tubes (357000, polycarbonate). The pellets were resuspended in 3 ml of microsome buffer; and aliquots were stored at -80^0^C in Eppendorf tubes.

##### Elongase activity assays

The elongase microsomal assay was performed in three sequential steps: biochemical reaction; hydrolysis and fatty acids extraction.

In ELOVL1 enzyme activity assay (1) 15ul of microsomes (75ug/ml, prepared in house) was preincubated with 15ul of substrate mixture containing 10uM d4C22-CoA(Avanti Polar Lipids, custom labeled), 350nM Malonyl-CoA(Sigma M4263), 250uM NADPH(Sigma N7785) at room temperature for 90 minutes in a 384-well Waters plate using a Hepes buffer (50mM Hepes pH6.8 (Sigma H0887), 5uM rotenone (Calbiochem 557368), 10uM BSA (Sigma A8806), 150mM NaCl2(Sigma S6546), 0.1mM MgCl2(Sigma M1028), 0.1mM CaCl2(Fluka 21115). (2) For the in-plate hydrolysis, 15uL of KOH (7.5N, Sigma 60370) was added to each well, plate was sealed, centrifuged, and samples were incubated at 80°C for 3 hours. (3) For fatty acid extraction, 50uL of acetonitrile (EMD AX0156) was added to each well, plate was sealed, and centrifuged at 3000 x g for one minute. Plates then were loaded onto the LX4-TSQ Quantiva system (X) and only the top layer of the samples were injected for MS measurement, monitoring the elongation of d4C22 to d4C24.

ELOVL3 biochemical assay was performed in analogous to ELOVL1 assay manner with several modifications: in step (1) – microsome concentration was 30ug/ml; substrate mixture was 7.5uM d4C18-CoA (Avanti Polar Lipids, custom labeled), 2.5uM Malonyl-CoA(Sigma) and incubation time was 60 min. In step (2) KOH concentration used was 0.5N and hydrolysis reaction was performed at room temperature for 30 min. In step (3) the elongation of d4C18 to d4C20 was monitored.

ELOVL7 enzyme activity assay was performed in analogous to ELOVL3 assay manner with exception: in step (1) 1μM malonyl CoA (Sigma M4263) was used in the substrate mixture.

##### Cell-based assay to study ELOVL1 inhibitors in human patients’ fibroblasts

Human patient fibroblasts isolated from a 6-year-old male Caucasian with elevated C26 VLCFA levels (Coriell Institute, CCALD4496) were plated to 384-well cell assay plates (Corning Cellbind, 3770) at 750 cells per well in 50 uL DMEM media (Invitrogen, #10566) with 10% FBS (Invitrogen, #16000-044) and 1% penicillin-streptomycin (Invitrogen, #15140); and incubated for 72 hours at 37°C in the absence or presence of compound dispensed in a 10-point dose response manner with 3-fold dilutions from 10 µM down to 0.5 nM, final concentrations (50 nL per concentration/well) using Beckman, ECHO655. After washing the plates with 1xPBS twice, residual PBS volume was removed, and the plates were dried in a biosafety cabinet. Plates were either heat-sealed and frozen at -20°C for future analysis or immediately underwent lipids extraction procedure: 85 uL of a 50:50 acetonitrile and methanol solution with 0.1 uM each of phosphatidylcholine (PC) C17:0 and sphingomyelin C17:0 standards included for normalization, were added to each well. The plates were centrifuged for 5 minutes at 3000 rpm and 75 uL of the cell extraction was transferred to a 384-well assay plate (Waters, 186002631) using the Apricot iPipette. The plates were heat-sealed (Agilent PlateLoc) and analyzed on the Rapidfire (Agilent) API5000 (Sciex) MS/MS system to determine sphingomyelin C26:0 to C22:0 ratio.

##### Data availability

Single-nuclei sequencing data of mouse study to be made available on GEO/SRA upon publication.

